# Resilience and restoration from fasting-refeeding mediated by a nutrient-regulated linker histone

**DOI:** 10.1101/2025.04.14.648802

**Authors:** Kazuto Kawamura, Anna R. Diederich, Birgit Gerisch, Roberto Ripa, Christian Latza, Joachim D. Steiner, Stephanie Fernandes, Filippo Artoni, David H. Meyer, Damini Sant, Simon Oehm, Franziska Grundmann, Roman-Ulrich Müller, Constantinos Demetriades, Adam Antebi

## Abstract

Intermittent fasting and fasting-refeeding regimens can slow biological aging across taxa^1^. Shifts between fed and fasted states activate ancient nutrient-sensing pathways which alter cellular and epigenetic states to promote longevity^2–4^. Yet how biological age trajectories progress during fasting-refeeding, and how nutrient-sensing pathways reprogram epigenetic state remain largely unknown. Here we observe increases in predicted biological age of *Caenorhabditis elegans* during prolonged fasting in adult reproductive diapause, followed by extraordinary reduction of biological age during refeeding. We identify *hil-1*/*H1-0* as an evolutionarily conserved nutrient-regulated linker histone which mediates adaptations to fasting and refeeding downstream of FOXO and TFEB transcription factors. In *C. elegans* and human cell culture, *hil-1*/*H1-0* upregulation during low-nutrient states promotes long-term survival and subsequent refeeding-induced recovery. Restoration of *C. elegans* after prolonged fasting is improved by enhancing the natural downregulation of *hil-1* specifically during refeeding. Our study identifies HIL-1/H1.0 as part of an ancestral epigenetic switch during fasting-refeeding that reprograms metabolic and cellular states underlying resilience and restoration.

## Main

Organisms respond vigorously to shifts in nutrient availability by adjusting their metabolism and development, to ensure long-term survival and reproductive capacity. This response is under strong evolutionary selection, and likely explains why dietary restriction paradigms and modulation of nutrient-sensing pathways are the most robust interventions to extend lifespan across taxa^5^. Both fasting and refeeding phases are likely to coordinate positive effects on organismal survival and function, but how they do so is not well understood.

### *C. elegans* age during adult reproductive diapause and transcriptomically rejuvenate upon refeeding

*C. elegans* can enter diapause states during its life cycle in response to food deprivation^6,7^. Developmental diapause has been described as a “non-aging state”^8,9^, yet other studies reveal that features of aging accrue during diapause, some of which are reversed upon exit^10,11^. *C. elegans* can enter adult reproductive diapause (ARD)^12^ in response to fasting at the mid-L3 stage and progress to mini-adults in a sleep-like quiescence^13^. Upon refeeding they restore their germline through re-proliferation and their soma by regrowth, emerging as young healthy adult worms with a normal lifespan (Fig. 1a)^12,13^. To quantitatively address whether adult diapause is an aging or non-aging state, we analyzed the transcriptomes of animals in prolonged ARD and compared them with transcriptomic changes associated with *ad libitum* (AL) aging^14^. During ARD, animals had an age trajectory that increased with time, as assessed by two independent transcriptomic clocks, BiT age^15^ and stochastic age^16^ (Fig. 1b; Extended Data Fig. 1a,b). Other transcriptomic changes during prolonged ARD also overlapped with changes during AL aging, including decreased reads aligned to protein-coding regions, increased snoRNA, antisense RNA, and transposable element (TE) expression (Extended Data Fig. 1c-j; Extended Data Table 1,2). Global gene expression changes accompanying ARD and AL aging positively correlated (r = 0.56) (Fig. 1c; Extended Data Table 3), and functional enrichment (Gene Ontology: Biological Process (GO:BP) and Kyoto Encyclopedia of Genes and Genomes (KEGG) pathway) suggested common aspects in their aging trajectories (Extended Data Fig. 1k,l; Extended Data Table 4). Importantly, ARD progression shared multiple features with AL aging, though ARD worms lived much longer.

**Fig. 1.**
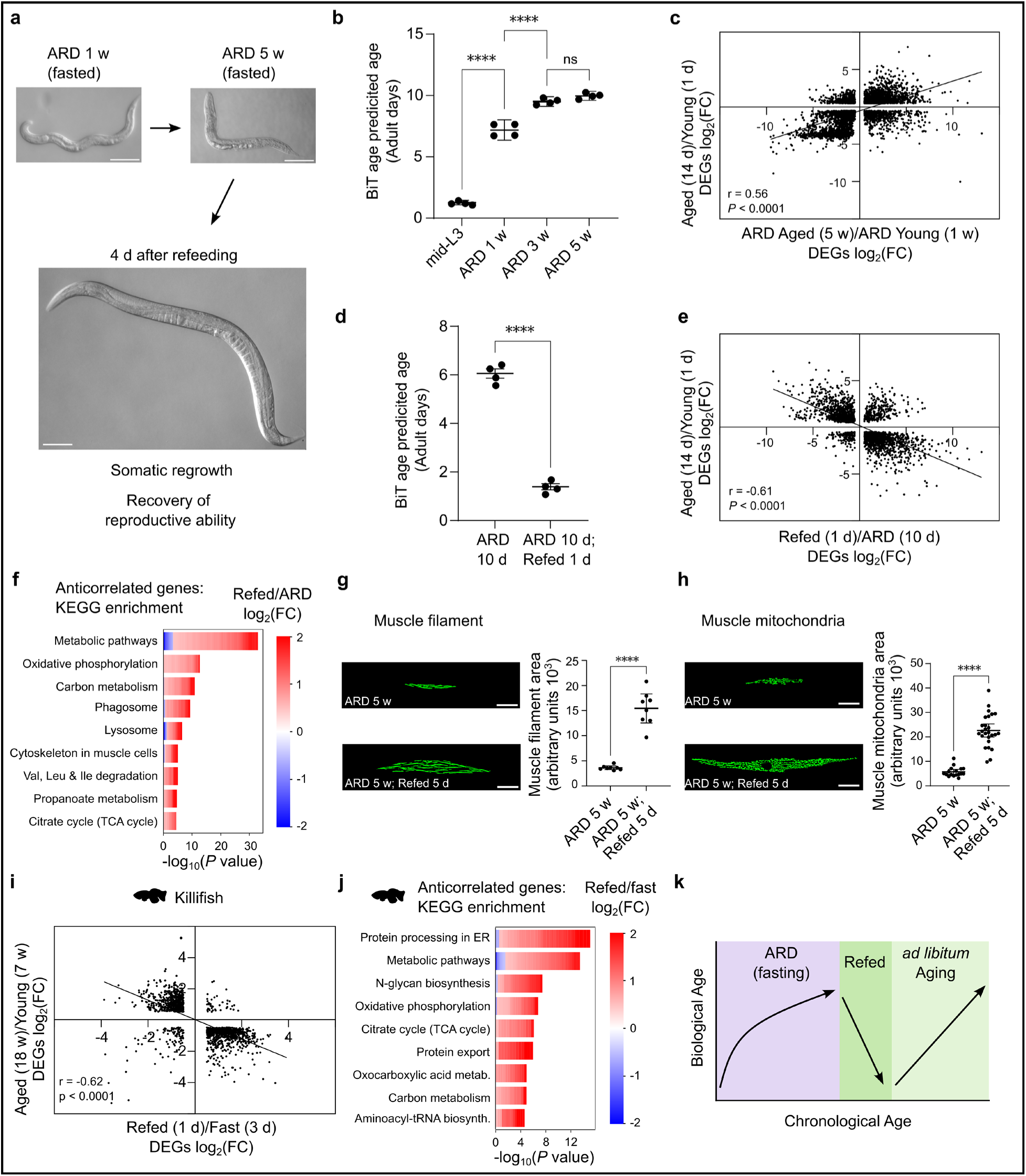
Progression and restoration of biological age during fasting-refeeding. **a**, Images of worms in ARD for 1 w, 5 w, and 4 d refeeding after 5 w of ARD. Scale bars 100 μm. **b**, BiT age biological age predictions based on transcriptomes from worms prior to ARD induction at mid-L3 stage, and ARD for 1, 3, and 5 w. Maximum biological age set at 15.5 days (See Methods – Age Prediction analysis for details). N = 4. One-way ANOVA with Tukey’s multiple comparisons test. **c**, Correlation plot of DEGs in AL aging (14 d vs. 1 d adult) and ARD aging (5 w vs. 1 w ARD). Adj. *P* < 0.05 log_2_(FC) > 0.5, < -0.5. **d**, BiT age biological age predictions of 10 d ARD and Refed (ARD 10 d followed by 1 d refeeding). Percentage of transcriptome reads aligned to protein coding regions from worms prior to ARD induction at mid-L3 stage, and ARD for 1, 3, and 5 w. N = 4. **e**, Correlation plot of DEGs in AL aging (14 d vs. 1 d adult) and refeeding from ARD (1 d Refed after 10 d ARD vs. 10 d ARD). Adj. *P* < 0.05 log_2_(FC) > 0.5, < - 0.5 and germline genes removed to rule out that the inverse correlation occurs solely from re-expression of germline-related genes during refeeding from ARD. See Methods for details. **f**, Top KEGG terms enriched in anticorrelated DEGs between AL aging and recovery from ARD. Red indicates upregulation during refeeding, blue indicates downregulation. *Adj. *P* < 0.05. **g**, Structure of muscle filaments (*unc-54::gfp*) and **h**, mitochondria (*mito::gfp*) in ARD (5 w) vs. recovery from ARD (Refed 3 d after ARD 5 w), with quantifications on right. N = 2. Scale bars 20 μm. **i**, Correlation plot of aging (18 w vs. 7 w) and refeeding (Refed 1 d after Fasting 3 d) from killifish visceral adipose. **j**, Top KEGG terms enriched in anticorrelated DEGs between AL aging and refeeding in killifish. **k**, Schematic of how shift from fasting to refeeding is a time window of biological age restoration. Unpaired t-test was used for comparisons between two conditions. Pearson correlation was calculated for comparing two transcriptomes with each other, and two-tailed *P* value was calculated to test whether the correlation was significantly non-zero. \*\*\*\**P* < 0.0001. Error bars represent mean ± SEM; for B, D; 95% confidence intervals for **g**, **h**.

During refeeding following ARD, remarkable reductions in predicted biological age occurred, suggesting transcriptomic rejuvenation (Fig. 1d, Extended Data Fig. 2a). Specifically, 10-day ARD worms displayed a transcriptomic age corresponding to 6.1 adult days AL, yet within a day of refeeding, showed reversal to 1.4 adult days (Fig. 1d). Refeeding also opposed general age-related transcriptome changes including increased reads aligned to protein-coding regions, and reduced intron retention, antisense RNA, snoRNA and TE expression (Extended Data Fig. 2b-f; Extended Data Table 5). Accordingly, a comparison of differentially expressed genes (DEGs) during refeeding and AL aging indicated a prominent inverse correlation (r = - 0.61), (Fig. 1e; Extended Data Fig. 2g; Extended Data Tables 6-8). Functional enrichment terms from the inversely correlated gene set suggested that refeeding partially reversed age-related changes related to metabolism, mitochondria and muscle function (Fig. 1f; Extended Data Fig. 2h,i; Extended Data Table 9). On the physiologic level, expansive regrowth of myofilaments and mitochondrial biogenesis accompanied refeeding (Fig. 1g,h)^17^. Rather than simply slowing down the aging process, these observations suggest a reversal in age-associated trajectories during refeeding.

To test whether similar trends could be seen in vertebrates, we analyzed fasted and refed transcriptomes in the turquoise killifish (*Nothobranchius furzeri*) visceral adipose, a tissue undergoing substantial changes during fasting-refeeding and aging^18,19^. Similar to *C. elegans*, transcriptomic changes upon refeeding inversely correlated with transcriptomic changes during aging (r = -0.65) (Fig. 1i; Extended Data Table 10). Functional enrichment analysis of the inversely correlated gene set identified terms related to metabolism and mitochondria, several of which overlap with *C. elegans* refeeding (Fig. 1j; Extended Data Fig. 2j,k; Extended Data Table 11). Our results indicate that both fasting and subsequent refeeding reprogram metabolic and cellular states, and refeeding resets biological age (Fig. 1k).

### *hil-1/H1-0* is an evolutionarily conserved nutrient-regulated linker histone

To identify functional mediators that facilitate restoration during the shift from fasting to refeeding, we carried out an RNAi screen for factors that impede or enhance recovery from ARD upon refeeding. We measured body size after refeeding, as it reflects the ability to re-access youthful genetic programs related to growth and the extent of restoration following diapause states^10^. For screening, we selected genes that were specifically up- or down-regulated during post-ARD refeeding, had a human ortholog, and had no reported phenotypes of lethality, sterility, or arrest, to rule out non-specific effects (Fig. 2a; Extended Data Table 12). We screened 380 genes by recovering worms after five weeks in ARD on RNAi-expressing bacteria and measured body size after three days (Fig. 2a,b). Thirty-six RNAi clones improved regrowth by over 15% vs. *luciferase* control (*luci* ctrl.) while 32 clones impeded regrowth by over 15% (Fig. 2b; Extended Data Fig. 3a; Extended Data Table 13). To factor in evolutionary conservation, we examined proteomic profiles of a human cell line (HEK293FT) in response to amino acid (AA) starvation, a well-established *in vitro* nutrient restriction paradigm^20^. AA starvation elicits adaptive cellular responses, in part via inhibition of the conserved mechanistic/mammalian target of rapamycin complex I (mTORC1), a master regulator of cellular growth and metabolism^20^. After cross-referencing (Extended Data Fig. 3a; Extended Data Table 14), the linker histone *hil-1/H1-0* stood out as the top candidate since its knockdown led to strong improvement of refeeding-induced regrowth (Fig. 2b) and its expression was strongly upregulated in both ARD (Fig. 2c; Extended Data Table 15) and HEK293FT AA starvation (Fig. 2d; Extended Data Table 16). Additionally, H1.0 was significantly upregulated in proteomic profiles of HEK293FT cells subjected to pharmacological mTOR inhibition (Fig. 2e; Extended Data Table 17) as well as killifish fasting-refeeding transcriptomics (Extended Data Fig. 3b; Extended Data Table 18)*. H1-0* encodes a linker histone that binds DNA regions between nucleosomes, maintaining chromatin compaction and epigenetic state^21–24^. Linker histones have been reported to modulate the transcriptome^25–28^, but its physiological function, especially in fasting-refeeding, is not known.

**Fig. 2.**
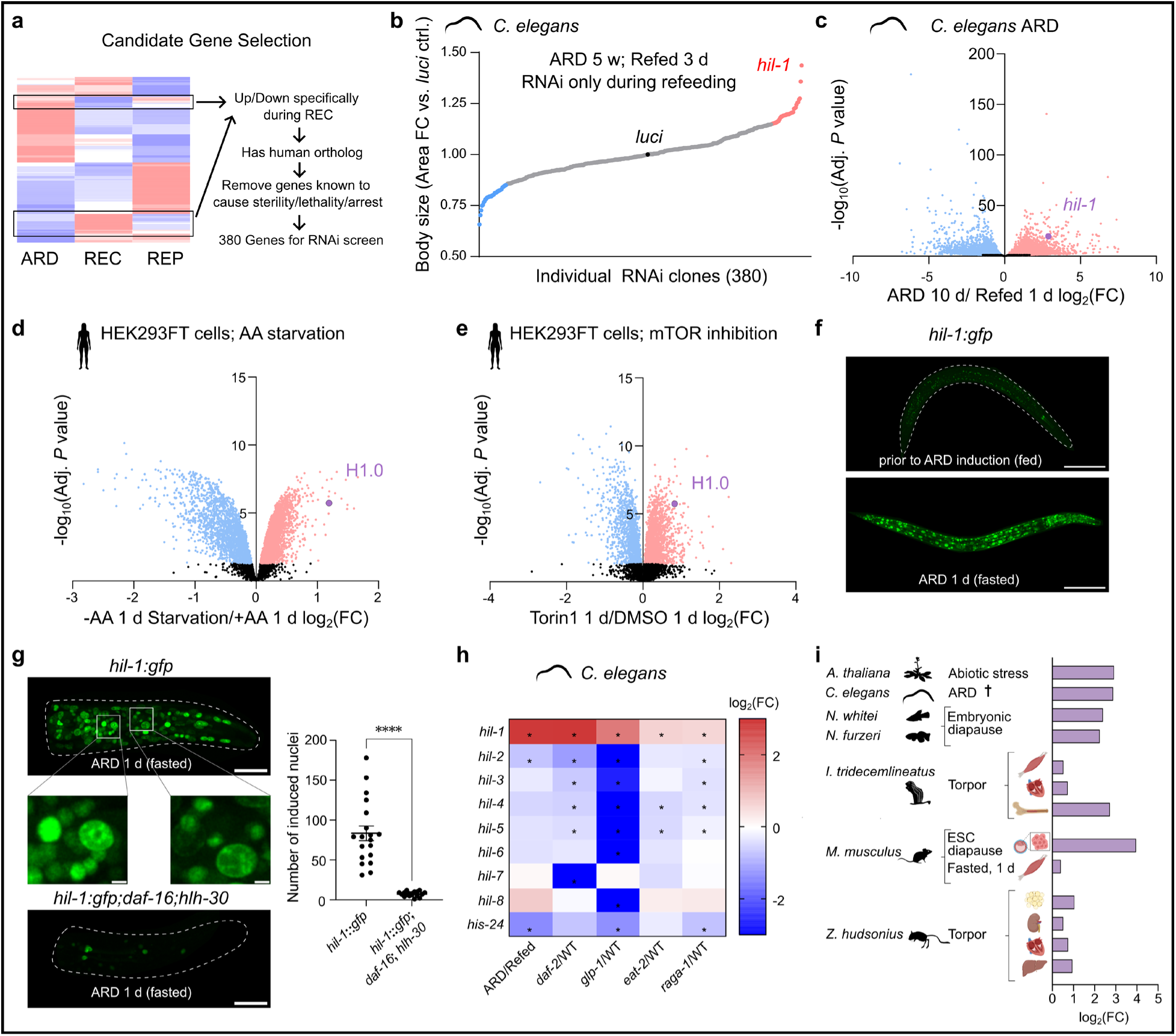
*hil-1/H1-0* is an evolutionarily conserved nutrient-regulated linker histone. **a**, Schematic of candidate gene selection process for ARD-refeeding RNAi screen. **b**, Fold change (FC) of body size area relative to *luci* ctrl. for individual RNAi knockdown of 380 genes. Enhancement of regrowth over 15% are colored red, inhibition of regrowth by over 15% are colored blue. Dots indicate the mean body size of surviving worms from 30 refed worms distributed over three plates. **c**, Volcano plot of DEGs during ARD 10 d vs. Refed 1 d. **d**, Volcano plot of differentially expressed proteins during -AA 1 d Starvation vs. +AA 1 d (AA-replete) conditions. **e**, Volcano plot of differentially expressed proteins during Torin1 (250 nM) 1 d treatment vs. DMSO 1 d control. **f**, Images of HIL-1::GFP expression prior to and after ARD (1 d) induction. Scale bar 100 μm. **g**, Head images of HIL-1::GFP expression in WT and *daf-16(mu86)*;*hlh-30(tm1978)* double mutant background. Scale bar 20 μm. Zoomed in images of some selected nuclei in middle panels. Zoomed in scale bar 2 μm. (Right) Quantification of number of head nuclei with HIL-1::GFP induction at ARD 1 d in WT and *daf-16(mu86)*;*hlh-30(tm1978)* worms. N = 3. Error bars represent mean ± SEM. **h**, Summary of transcriptomic regulation of *C. elegans* linker histones during ARD and longevity contexts (*daf-2(e1370)*, *glp-1(e2141)*, *eat-2(ad465), raga-1(ok386)*). Significantly up- or downregulated genes (Adj. *P* < 0.05) are indicated with *. **i**, Summary of statistically significant *hil-1*/*H1*-*0* regulation in low nutrient conditions across organisms from both publicly available transcriptomics data and our RNA-sequencing analysis (marked with **^†^**). For *A. thaliana*, *HIS1-3*, a drought-induced linker histone with similarity to *H1-0* is shown. Unpaired t-test \*\*\*\**P* < 0.0001. In volcano plots, significantly upregulated genes/proteins (Adj. *P* < 0.05) are colored red; significantly downregulated genes/proteins are colored blue.

To determine the expression pattern of *hil-1* in *C. elegans*, we created a knock-in strain tagging endogenous *hil-1* with *gfp*. During ARD, HIL-1 was strongly induced in somatic nuclei, but not the germline (Fig. 2f). In some cases, punctate HIL-1 expression or enrichment at the nuclear and nucleolar periphery occurred, reminiscent of fasting-induced chromatin reorganization (Fig. 2g)^29,30^. In AL conditions, *hil-1* was weakly expressed in a few somatic tissues, namely marginal cells, spermathecal-uterine valve, and vulval muscle cells (Fig. 2f; Extended Data Fig. 3c), overlapping with a previous report of HIL-1 immunostaining^31^. Notably, we found that upregulation of *hil-1* during ARD was dependent on nutrient-responsive transcription factors (TFs) DAF-16/FOXO and HLH-30/TFEB (Fig. 2g), key regulators of ARD survival as well as canonical longevity pathways^13,32,33^. This is likely to be the result of a direct interaction between these two TFs and the *hil-1* promoter, which is consistent with existing ChIP-Seq data^32^.

Intriguingly, *hil-1* expression was also upregulated in several longevity pathways, including in *daf-2(e1370)* (insulin/IGF signaling), *glp-1(e2141)* (reproductive signaling), *eat-2(ad465)* (dietary restriction), and *raga-1(ok386)* (mTOR signaling) mutants under AL conditions (Fig. 2h). *hil-1* was the only linker histone significantly upregulated in both ARD and longevity contexts, suggesting its potential subtype-specific role. Further mining of existent transcriptomics data indicated that *H1-0* was upregulated in other hypometabolic states across species, including fasting, torpor, embryonic diapause and *in vitro* embryonic stem cell diapause (Fig. 2i)^34–41^. Remarkably, even in *Arabidopsis thaliana*, a plant linker histone *HIS1-3* is upregulated in response to drought (Fig. 2i)^42^. Altogether, these consistent changes suggest that upregulation of *hil-1*/*H1-0* is an evolutionarily ancient response to low-nutrient conditions.

### *hil-1* promotes longevity and a fasting-adapted transcriptome

Since *hil-1* expression was upregulated during ARD and in various longevity pathways, we tested the effect of *hil-1* mutation on lifespan. In ARD, *hil-1* mutants were significantly shorter-lived (∼20%) compared to WT (Fig. 3a), though AL lifespan was unchanged (Extended Data Fig. 4a). Accordingly, *hil-1(tm1442)* mutants had older predicted biological ages after prolonged ARD (Extended Data Fig. 4b). *hil-1(tm1442)* was also partly required for prolonged lifespan in *eat-2(ad465)* (Fig. 3b) and *glp-1(e2141)* (Fig. 3c), but not *daf-2(e1370)* mutants (Extended Data Fig. 4c). Thus, *hil-1* promotes longevity in several contexts.

**Fig. 3.**
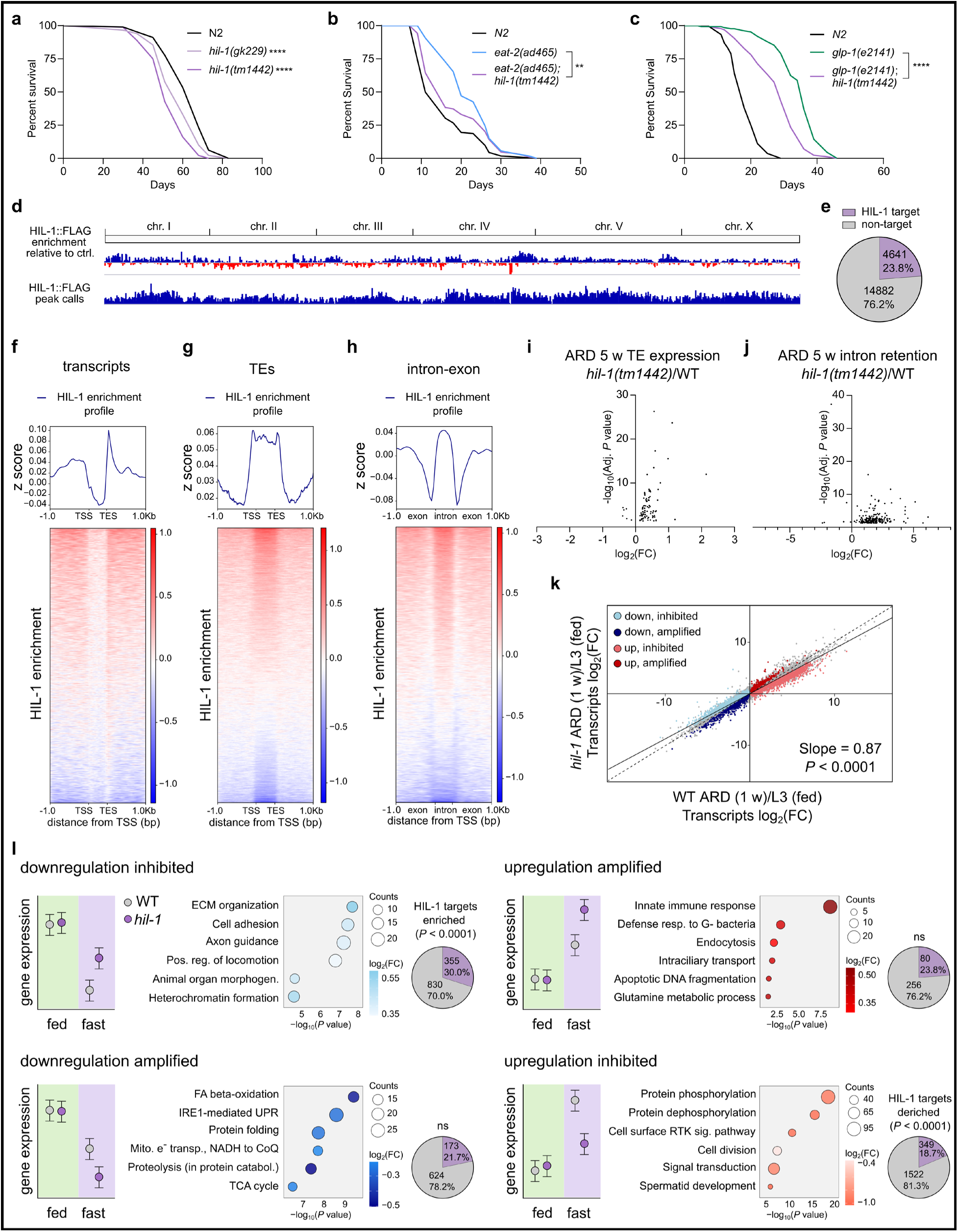
HIL-1 directly affects the transcriptome for fasting adaptation. **a,** Lifespan of WT, *hil-1*(*tm1442*), *hil-1*(*gk229*) worms in ARD. N = 4. **b**, Lifespan of *eat-2(ad465)* dietary restriction model compared to *eat-2(ad465)*; *hil-1(tm1442)* double mutants. N = 4. **c**, Lifespan of *glp-1(e2141)* germline-less longevity model compared to *glp-1(e2141);hil-1(tm1442)* double mutants. N = 3. **d,** (Upper panel) HIL-1:FLAG enrichment relative to FLAG antibody control at 1 w ARD. (Lower panel) Binding peaks of HIL-1::FLAG, from MACS3 peak caller at 1 w ARD. Pooled from N = 4. **e**, Proportion of HIL-1 target genes relative to total genes. **f**, Enrichment of HIL-1:FLAG binding relative to transcription start and end sites for transcripts, and **g**, TEs, and **h**, intron-exon boundaries. **i**, Volcano plots of significantly differentially expressed TEs and **j**, intron reads at ARD 5 w in WT vs. *hil-1(tm1442)* mutants. (Adj. *P* < 0.05). **k**, Transcriptomic comparison of log_2_(FC) of WT response to ARD induction 1 w vs. *hil-1(tm1442)* mutant response to ARD induction 1 w. Significant DEGs from the genotype-ARD interaction term are highlighted in colors according to category. Nonlinear fit analysis compared the observed slope against a hypothetical correlation of x = y, Slope = 1. *P* < 0.0001. **l**, DEGs from the genotype-ARD interaction term, divided into four categories: ‘up inhibited’, ‘up amplified’, ‘down inhibited’, ‘down amplified’. Top 6 enriched GO:BP terms are displayed next to each category. Interaction log_2_(FCs) are indicated in color, with darker colors indicating greater inhibited or amplified responses in the *hil-1* mutant. Circle sizes are scaled to counts, which are the number of genes in each category. Number of HIL-1 targets from CUT&RUN that overlap with DEGs in each category are displayed in the pie chart. Whether the overlap is significantly enriched or deriched compared to expected probability (**e**) was tested by the Binomial test. Statistical significance between lifespan curves was tested with log-rank (Mantel-Cox) test. *P*\*\* < 0.01; *P*\*\*\*\* < 0.0001.

To identify the direct genomic targets of *hil-1* during ARD, we performed cleavage under target and release under nuclease (CUT&RUN) sequencing^43^. We identified 8058 putative HIL-1 binding sites, 4641 of which resided in a gene (including the promoter and 3’UTR) (Fig. 3d,e; Extended Data Tables 19,20). HIL-1 was enriched near transcription start sites and moreso near transcription end sites while deriched in gene bodies (Fig. 3f). By contrast, HIL-1 was enriched throughout the entire sequence of TEs (Fig. 3g). HIL-1 was also differentially bound to introns versus exons, hinting at a possible role in directing the splicing machinery (Fig. 3h). The direct targets of HIL-1 overlapped most with targets of PHA-4/FOXA1 and BLMP-1/BLIMP (Extended Data Fig. 4d; Extended Data Table 21), two transcription factors known to mediate dietary restriction-induced longevity^44,45^. HIL-1 targets also significantly overlapped with those of DAF-16/FOXO (*P* = 3.2×10^-10^) (Extended Data Table 21), which itself regulates HIL-1 (Fig. 2d), suggesting co-regulation between DAF-16 and HIL-1.

To understand if HIL-1 modulates the fasted transcriptome, we performed RNA-sequencing of *hil-1(tm1442)* mutant worms in mid-L3 (fed) and ARD (fasted) (Extended Data Fig. 4e)*. hil-1(tm1442)* mutants showed a strong skew towards increased TE expression and intron retention in ARD, but not in the fed condition (Fig. 3i,j; Extended Data Fig. 4f,g; Extended Data Table 22,23). We also measured transcriptional speed, using the slope of intronic reads as a proxy^46^, and found that it was slower in ARD compared to AL, and further slowed down during ARD in the *hil-1* mutant (Extended Data Fig. 4h), suggesting that HIL-1 is required for optimal transcriptional elongation. Linear regression analysis comparing *hil-1* and WT transcriptomes showed a slope < 1 (0.87), suggesting that *hil-1* mutation blunts the fasting response, and that it normally promotes adaptation to the fasted state by influencing both gene activation and repression (Fig. 3k). We carried out a genotype-diet interaction analysis^47^ to identify fasting-responsive genes that were *hil-1*-dependently dysregulated and grouped these genes based on directionality of change during starvation (upregulated or downregulated) and effect of mutation on this response (inhibited or amplified) (Fig. 3l; Extended Data Table 24). The ‘down inhibited’ category, encompassing genes for which *hil-1* directly or indirectly plays a role in their repression, overlapped most with HIL-1 targets (Fig. 3l). This is consistent with the role of linker histones in gene repression^21,24^. Functional enrichment terms in the ‘down inhibited’ category showed derepression of genes related to morphogenesis, development and heterochromatin organization as well as metabolic genes in the *hil-1* mutant (Fig. 3l; Extended Data Fig. 4i; Extended Data Table 25). The ‘up inhibited’ category suggested *hil-1* mutants had suboptimal induction of genes related to DNA damage repair, polycomb repressive complex and longevity-regulating pathways (Fig. 3l; Extended Data Fig. 4i; Extended Data Table 25). Taken together, our findings suggest that HIL-1 promotes a fasting-adapted transcriptome by optimizing transcriptional speed, preventing transposable element expression and developmental programs, as well as facilitating intron removal and longevity programs.

### *hil-1* regulation is required for optimal recovery from adult reproductive diapause

Since *hil-1* mutants were maladapted during ARD and *hil-1* expression was strongly downregulated upon refeeding (Fig. 4a), we wondered if *hil-1* mutation would affect refeeding-induced recovery. *hil-1* mutants in AL or refed from short-term ARD (1 w) were only slightly (∼10%) smaller than WT (Extended Data Fig. 5a,b), but substantially (40.7%) smaller when refed after long-term ARD (5 w) (Fig. 4b). After long-term ARD and refeeding, *hil-1* mutants also took longer to regain reproductive ability, had considerably smaller brood size (Extended Data Fig. 5c,d) and showed older transcriptional age compared to WT (Fig. 4c). Linear regression analysis comparing WT and *hil-1* transcriptomes during ARD recovery showed a slope < 1 (0.64), suggesting that *hil-1* mutation blunts the transcriptional response to refeeding (Fig. 4d; Extended Data Fig. 5e; Extended Data Table 26). Enriched terms in the ‘up inhibited’ category suggested that *hil-1* mutants had dampened chromatin remodeling and cell cycle reactivation, indicative of impeded somatic and reproductive recovery (Fig. 4e; Extended Data Fig. 5f; Extended Data Table 27). The ‘down inhibited’ category suggested an inability to completely shift neuropeptide signaling and turn off programs related to sleep (Fig. 4e; Extended Data Fig. 5f, Extended Data Table 27). We also noticed that HIL-1::GFP animals kept long-term in ARD displayed variable levels of expression prior to recovery. When animals were grouped by high or low expression level, body size recovery was significantly greater in animals expressing high levels prior to recovery (Fig. 4f). Hence, *hil-1* induction during ARD promotes survival and preserves restorative potential during long-term fasting.

**Fig. 4.**
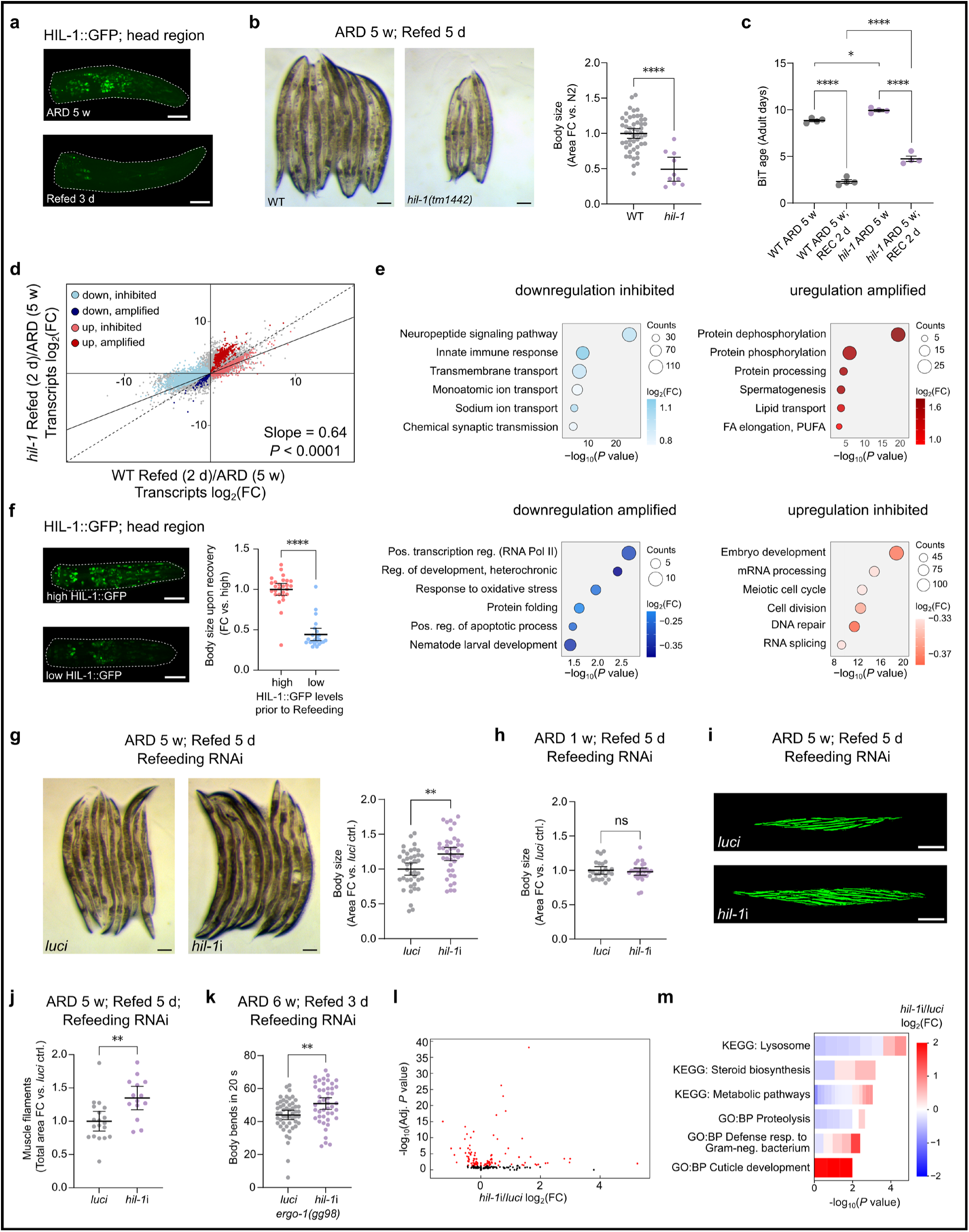
HIL-1 regulation is critical for refeeding-induced recovery. **a**, Images of HIL-1::GFP in WT ARD 5 w and Refed 3 d. **b**, Images of worms Refed 5 d after ARD 5 w for WT and *hil-1(tm1442)* worms, with quantifications on the right panel. N = 3. **c**, BiT age biological age predictions based on transcriptomes from worms at ARD 5 w and Refed 2 d in WT and *hil-1(tm1442)* mutant worms. **d**, Transcriptomic comparison of log_2_(FC) of WT response to refeeding vs. *hil-1(tm1442)* mutant response to refeeding. Significant DEGs from the genotype-refeeding interaction term are highlighted in colors according to category. Nonlinear fit analysis compared the observed correlation against a hypothetical correlation of x = y, Slope = 1. *P* < 0.0001. **e**, Top 6 enriched GO:BP terms of significant genes from the genotype-refeeding interaction analysis are displayed for each category. Interaction log_2_(FCs) are indicated in color, with darker colors indicating greater inhibited or amplified responses in the *hil-1* mutant. Circle sizes are scaled to counts, which are the number of genes in each category. **f**, Worms were split into high vs. low HIL-1::GFP prior to refeeding. Body size quantifications of worms Refed 5 d after ARD 5 w. N= 3. **g**, Images of worms Refed 5 d after ARD 5 w on *luci* ctrl. control vs. *hil-1* RNAi. Quantification on right. N = 5. **h**, Body size quantifications of worms Refed 5 d after ARD 1 w for WT worms on *luci* ctrl. or *hil-1* RNAi. N = 3. Images (**i**) and quantification (**j**) of muscle filaments of worms Refed 3 d after ARD 5 w on *luci* ctrl. or *hil-1* RNAi. N = 2. **k**, Body bending assay of worms Refed 3 d after ARD 6 w on *luci* ctrl. or *hil-1* RNAi. Experiments carried out in RNAi sensitive strain *ergo-1*(*gg98*). N = 3. **l**, Volcano plot of DEGs in worms Refed 3 d after ARD 5 w on *luci* ctrl. or *hil-1* RNAi. Red dots indicate significance (Adj. *P* < 0.05). **m**, Top 3 enriched GO:BP and KEGG terms each from DEGs in the *hil-1* RNAi condition (**l**). Red indicates upregulation by *hil-1* RNAi, whereas blue indicates downregulation. Error bars represent 95% confidence intervals, except for **c**, which is mean ± SEM. Statistical significance between two groups were tested by unpaired t-test or one-way ANOVA with Tukey’s multiple comparisons test for multiple comparisons. *P*\* < 0.05; *P*\*\* < 0.01; *P\*\*\*\** < 0.0001. ns = not significant.

Although *hil-1* mutants kept in ARD recovered poorly upon refeeding (Fig. 4b), we initially found and later confirmed that *hil-1* knockdown specifically during refeeding enhanced body size restoration following long-term ARD (Fig. 2b, 4g). *hil-1* knockdown during refeeding had no impact on body size recovery after short-term ARD (Fig. 4h). This could reflect a loss of plasticity whereby persistent fasting-related programs from long-term ARD impair restoration from refeeding. *hil-1* knockdown after long-term ARD not only enhanced body size, but also muscle regrowth (Fig. 4i,j). Using an RNAi-sensitive strain (*ergo-1*(*gg98*)) to enhance knockdown throughout the body, *hil-1* RNAi administered during refeeding after long-term ARD (6 w) also improved motility (Fig. 4k). Transcriptome analysis of worms refed after long-term ARD in *hil-1* and *luci* ctrl. RNAi conditions revealed 106 DEGs (*p*. adj < 0.05) (Fig. 4l; Extended Data Fig. 5g; Extended Data Table 28). Functional enrichment indicated that *hil-1* knockdown primarily affected metabolic processes (Fig. 4m; Extended Data Table 29). Interestingly, *hil-1* knockdown generally augmented the up- or downregulation of genes during recovery from ARD, in line with promoting the normative recovery process (Extended Data Fig. 5h). In sum, *hil-1* acts in a nutrient-regulated switch between fasting and feeding states–it is induced by fasting and promotes resilience during long-term diapause, while its downregulation during refeeding facilitates recovery. Notably, increasing the sharpness of this switch enhances restorative genetic programs.

### *H1-0* promotes adaptation to amino acid starvation and addback responses in human cell culture

To further decipher the potential evolutionary conservation of HIL-1/H1.0 regulation and function during nutrient shifts in human cell culture, we first validated H1.0 induction in response to low-nutrient states (Fig. 2e,f) by immunoblotting. H1.0 upregulation was confirmed in HEK293FT cells by 1 d Torin1 treatment and AA starvation (Fig. 5a), where it was visibly localized to the nucleus (Fig. 5b). In addition to H1.0, the subtype H1.2 displayed similar regulation, albeit to a lesser extent than H1.0 upon AA starvation (Extended Data Fig. 6a-d). Upregulation of H1.0 by both treatments was also seen in human transformed lung fibroblast WI-26 cells (Extended Data Fig. 6e). As in the worm, *H1-0* induction in HEK293FT cells was reversible upon addback after AA starvation (Extended Data Fig. 6f) and regulation occurred at the transcriptional level (Extended Data Fig. 6g,h). Combinatorial knockdown of *FOXO1* and *FOXO3* attenuated *H1-0* induction during AA deprivation (Figure S6i,j), suggesting conserved regulation through DAF-16/FOXO transcription factors. Accordingly, available ChIP-seq data revealed FOXO1 and FOXO3 binding in the *H1-0* promoter region in two different cell lines^48^ (GSE80773, GSE97661) (Extended Data Fig. 6k).

**Fig. 5.**
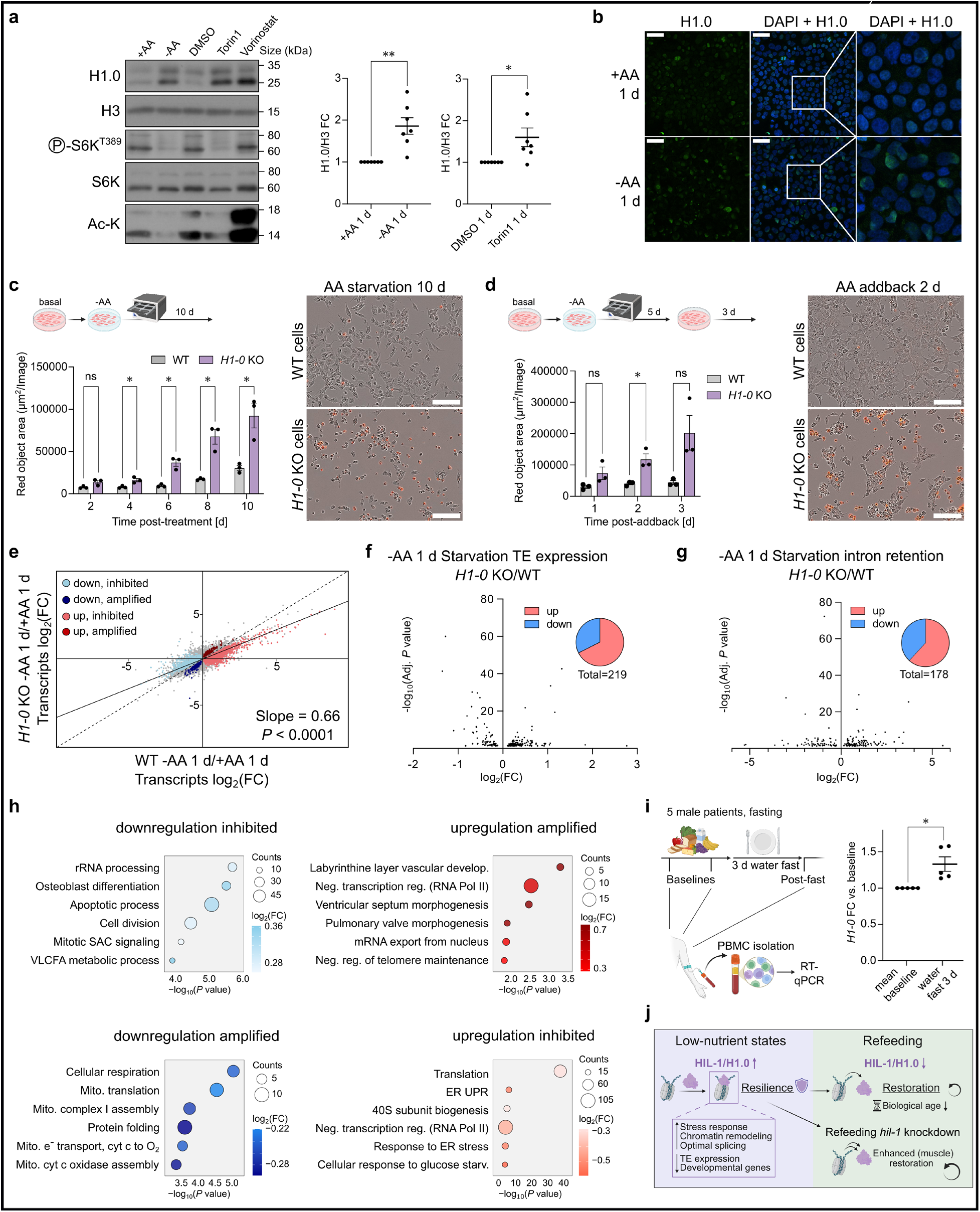
*H1-0* is a functional regulator of amino acid starvation in human cell culture. **a**, Immunoblots of lysates from HEK293FT cells under -AA 1 d Starvation vs. +AA 1 d conditions and Torin1 1 d (250 nM) vs. DMSO 1 d control, probed with the indicated antibodies. The histone deacetylase inhibitor Vorinostat (10 µM) was used as a control for H1.0 induction ^60^. N = 3. **b**, Immunofluoresence staining of H1.0 in HEK293FT cells under -AA 1 d Starvation vs. +AA 1 d conditions. Scale bar, 50 µm; N = 3. **c**, Evaluation of cell death in HEK293FT *H1-0* KO cells during -AA 10 d Starvation, as measured by the Incucyte Cytotoxicity Assay. Cell death was quantified by Red object area. N = 3. Representative images after -AA 10 d Starvation are shown to the right; scale bar, 150 µm. **d**, Evaluation of cell death in HEK293FT *H1-0* KO cells upon AA addback after -AA 5 d Starvation, as measured by the Incucyte Cytotoxicity Assay. Cell death was quantified by Red object area. N = 3. Representative images of AA addback 2 d are shown to the right; scale bar, 150 µm. **e**, Transcriptomic comparison of log_2_(FC) of WT response to -AA 1 d Starvation vs. *H1-0* KO response to -AA 1 d Starvation. Significant DEGs from the genotype-starvation interaction term are highlighted in colors according to category. Nonlinear fit analysis compared the observed correlation against a hypothetical correlation of x = y, Slope = 1. *P* < 0.0001. **f**, Volcano plot of significantly (Adj. *P* < 0.05) differentially expressed TEs after -AA 1 d Starvation in WT vs. *H1-0* KO cells. **g**, Volcano plot of significantly (Adj. *P* < 0.05) differentially expressed intron reads after -AA 1 d Starvation in WT vs. *H1-0* KO cells. **h**, Top 6 enriched GO:BP terms of significant genes from the genotype-starvation interaction analysis are displayed for each category. **i**, *H1-0* mRNA levels were assessed by RT-qPCR in PBMCs isolated from five patients with autosomal dominant polycystic kidney disease (ADPKD) after a 3 d water fast. *H1-0* expression FC was calculated relative to the mean of two AL baseline values. Ribosomal Protein Lateral Stalk Subunit P0 (*RPLP0*) was used as a housekeeping gene. **j**, Graphical summary of HIL-1/H1.0 regulation and function under low-nutrient states and refeeding. Error bars represent mean ± SEM. Statistical significance between two groups was tested by unpaired one-sample t-test (comparing against a hypothetical value of one) while the Incucyte timecourse was assessed by multiple unpaired t tests with Holm-Šídák correction. ns = not significant. *P*\* < 0.05; *P*\*\* < 0.01; *P\*\*\*\** < 0.0001. ns = not significant.

To evaluate the physiological function of *H1-0*, we generated *H1-0* knockout (KO) HEK293FT cells (Extended Data Fig. 7a). Under basal nutrient-replete culture conditions, *H1-0* KO cells proliferated at a similar pace as control cells (Extended Data Fig. 7b). However, during long-term (10 d) AA starvation, *H1-0* KO cells underwent increased cell death from day 4 onwards (Fig. 5c). Additionally, *H1-0* KO cells underwent increased cell death upon AA addback after prolonged (5 d) but not short-term (1 d) AA starvation, concomitant with a decrease in cell confluency (Fig. 5d, Extended Data Fig. 7c-e), suggesting that *H1-0* is required for cellular resilience, analogous to worm HIL-1.

To determine the effects of *H1-0* mutation on the fasting response, we performed transcriptomic analysis of HEK293FT *H1-0* KO cells and WT cells under AA-replete and 1 d AA starvation conditions (Extended Data Fig. 7f; Extended Data Table 30). Linear regression analysis comparing the transcriptome of *H1-0* KO cells to WT cells showed a slope < 1 (0.66), suggesting attenuation of the starvation response (Fig. 5e). Also similar to *C. elegans hil-1* mutants, TEs were derepressed and intron retention events increased under starvation conditions (Fig. 5f,g; Extended Data Tables 31,32). GO:BP and KEGG pathway enrichment analysis showed that rRNA processing was derepressed while translation was dampened, suggesting dysregulation of protein synthesis in *H1-0* KO cells during AA starvation (Fig. 5h; Extended Data Fig. 7g; Extended Data Table 33). Induction of transcriptional regulators and stress response genes was inhibited, including ATF4, a key regulator of the amino acid response (AAR) pathway^49^. ATF4 targets such as ATF3 along with several AA transport and metabolism genes were among the most significantly regulated genes in the ‘up inhibited’ category, suggesting blunting of the AAR pathway in *H1-0* KO cells. Concurrently, transcripts associated with development and cancer were derepressed or further upregulated (Fig. 5h; Extended Data Fig. 7g; Extended Data Table 33), indicating that *H1-0* KO cells fail to activate resilience mechanisms and aberrantly induce growth-related programs during AA starvation, despite mTORC1 being inactivated.

GO:BP terms from *hil-1* ARD worms and AA-deprived *H1-0* KO cells overlapped significantly higher than expected by chance in the ‘up inhibited’ and ‘down amplified’ categories (Extended Data Table 34). Notably, genes involved in FOXO signaling and chromatin reorganization were blunted, while genes related to mitochondrial function were further downregulated (Extended Data Fig. 7h). Altogether, our data suggests that HIL-1/H1.0 promotes resilience during low-nutrient states in *C. elegans* and human cells through overlapping mechanisms.

Finally, to test whether *H1-0* responds to dietary interventions in humans, we carried out real-time quantitative PCR (RT-qPCR) of *H1-0* in peripheral blood mononuclear cells (PBMCs) from three-day fasted male patients compared to the mean of two baseline values before fasting. *H1-0* was significantly upregulated in all five patients to varying degrees (Fig. 5i). Accordingly, available transcriptomics data indicated *H1-0* was significantly induced in human PBMCs of one-day fasted healthy individuals, calorically restricted participants of the CALERIE study and cancer patients on a fasting-mimicking diet^50–52^ (Extended Data Fig. 7i). Additionally, H1.0 was significantly upregulated in naive human pluripotent stem cells induced into dormancy by mTOR inhibition (Extended Data Fig. 7i), and downregulated during release from mTOR inhibition^53^.

## Discussion

Our study suggests that aging is slowed down by coordinate resilience and restoration mechanisms. Through characterization of ARD, we identify fasting as a period of resilience, and refeeding as a remarkable period of organismal biological age restoration. Similar to how the genetic dissection of *C. elegans* dauer diapause opened up the field of molecular genetics of longevity, we believe refeeding from ARD represents a paradigm to dissect genetic and metabolic mechanisms of adult organismal restoration and rejuvenation.

*hil-1*/*H1-0* upregulation is an evolutionarily conserved resilience response to low-nutrient status, and acts downstream of key nutrient signaling pathways implicated in longevity such as mTOR and insulin signaling. Induction of *hil-1/H1-0* during fasting promotes survival and preserves long-term restorative potential upon refeeding, suggesting it may be especially important for maintaining prolonged quiescent states such as diapause, torpor, or quiescent stem cell populations. Dysregulation of *hil-1/H1-0* may contribute to age-related loss of quiescence or aberrant quiescent states such as a constitutive fasting-like transcriptional program in aged killifish adipose or hyper-quiescent chromatin in aged mouse tissues^18,54^. Interestingly, both *H1-0* and *H1-2* emerged as key components of single cell transcriptomic aging clocks developed from neurogenic regions of the brain^55^, suggesting they could be causal biomarkers reflecting increased quiescence during aging. *hil-1*/*H1-0* downregulation occurs in response to refeeding, and enhancing the natural downregulation of *hil-1* upon refeeding improves restoration of the *C. elegans* muscular system. During another state of biological age reduction, iPS cell reprogramming and partial reprogramming, *H1-0* expression is reduced^56–58^, analogous to regulation that occurs during the restorative refeeding phase. Enhancement of restoration by leveraging refeeding-induced changes may open new possibilities for reprogramming healthy longevity.

Metabolic state can influence chromatin state. It does so by altering the availability of small molecules, such as S-adenosyl-methionine, acetate, butyrate and NAD+, which are substrates or co-factors used to post-translationally modify core histones, and enumerate the epigenetic code^59^. In this work, we describe a novel parallel mechanism for how nutrient signaling can modify the epigenome–through regulation of a linker histone*. hil-1/H1-0* likely mediates the alignment of metabolic and epigenetic states during nutrient shifts by facilitating correct splicing, inducing stress response genes, and adapting chromatin while suppressing aberrant transcription of TEs and developmental genes. Future efforts may be directed at deciphering the linker histone code in contexts of resilience and restoration, and how it contributes to epigenetic changes during aging.

## Methods

### *C. elegans* strains and maintenance

*C. elegans* were maintained at 20°C following standard procedures^61^. N2 strain was used as WT. All strains used in this study are listed in Extended Data Table 35.

### Induction of worms into adult reproductive diapause

Adult reproductive diapause (ARD) was induced as previously described by starving worms at the mid-L3 stage^13^. The mid-L3 stage was determined as the time-point when the distal tip cell of the somatic gonad turns from the ventral to dorsal side of the worm. Worms were collected by washing off the plate with M9 buffer, waiting for the worms to settle to the bottom of a 5 mL eppendorf tube and replacing the supernatant with fresh M9 buffer. M9 buffer was replaced after 20 minutes, for a total of 3 washes following collection. ARD worms were maintained on Nematode Growth Medium (NGM) plates with UltraPureTM agarose instead of agar, and with 50 μg/ml ampicillin. Worms were maintained at 20°C.

### Recovery from ARD and body size measurements

ARD worms were recovered by gentle transfer using a flattened worm pick onto plates with bacteria (OP50 or HT115 RNAi bacteria). Worms were transferred onto an area of the plate where there was no bacteria, and checked until they started moving towards the food. Worms that did not move following the transfer were assumed to be unhealthy or dead and removed from the plate. Equal numbers of worms were recovered for each condition in an experiment. Body size of recovered worms was measured by anesthetizing worms with 50 mM sodium azide, lining them up and taking a photo with Leica M80 binocular microscope with Leica MC180 camera. Photos were captured using LAS X or LAS EZ software and analyzed by outlining the body of the worm with the polygon selection tool in ImageJ.

### Lifespan measurements

*Ad libitum* lifespans were carried out on a synchronized population of 100-125 worms per genotype, equally distributed over 4-5 plates. L4 stage was considered as Day 0. Worms were transferred every 2-3 days during the reproductive period, and scored for survival by a tap to the head and tail with a wormpick. Worms that did not respond were considered dead. Worms that died from egg-laying deficiency, ruptured vulva, or crawled off the plate were censored. ARD lifespans were carried out on a synchronized population of a few hundred worms per genotype distributed over 3-4 plates. Worms that did not respond to a gentle tap to the head and tail were considered dead and removed from the plate. ARD worms were not transferred to new plates over the course of the lifespan. Some WT N2 worm strains are known to have a *fln-2* mutation which affects *eat-2* longevity. In all strains used for lifespan comparisons, the *fln-2* allele was checked and made sure to be the same across compared conditions. Summary of lifespan measurements including median lifespan, total number of worms, and *P* values of all replicates are shown in Extended Data Table 36.

### RNA-sequencing and transcriptome analysis

RNA extraction was carried out by RNeasy Mini kit (cat. no. 74106, Quiagen). Library preparation was carried out after rRNA was depleted using Illumina Stranded Total RNA Prep with Ribo-Zero Plus (Illumina). 40 million paired end reads were sequenced per sample on Illumina HiSeq at the Cologne Center for Genomics (https://ccg.uni-koeln.de). RNA-seq reads were analyzed through a standardized pipeline established by the MPI-AGE bioinformatics facility. RNA-seq reads were aligned to the ce11 reference genome using kallisto^62^, and pair-wise differential gene expression was calculated using DESeq2^47^. Functional enrichment analysis was done by DAVID for Gene Ontology: Biological Process (GO:BP) and Kyoto Encyclopedia of Genes and Genomes (KEGG) pathway terms. In some cases, redundant terms were omitted for visualization, though full lists of all terms are available as Extended Data Tables. Correlation analyses were done with Graphpad Prism using correlation to calculate Pearson r coefficient, and nonlinear fit analysis to compare the observed correlation with a hypothetical correlation of x = y, Slope = 1. Germline genes were removed for correlation analyses for comparison between ARD and refed conditions (Fig. 1E, Extended Data Fig. 2G) based on germline-expressed genes detected from Wang et al.^63^ to rule out whether the inverse correlation occurred solely from re-expression of germline-expressed genes. Germline genes in Wang et al. were determined by expression analysis following germline dissection by cutting near the spermatheca. Therefore sperm genes may not be completely included in this list. For the genotype-diet interaction analysis, see “Transcriptome interaction analysis” below. Transposable element (TE) expression was analyzed using TEtranscripts package^64^. Intron retention was analyzed using SAJR splicing analysis package ^65^.

### Age prediction analysis

BiT age analysis was carried out based on source code from^15^ https://github.com/Meyer-DH/AgingClock. Counts-per-million normalized RNA-seq reads for each genotype were used as input for BiT age analysis. Unlike traditional clocks that estimate chronological age or require age acceleration calculations, BiT age directly predicts biological age based on binarized transcript expression of 576 genes. A maximum biological age of 15.5 days was set based on lifespan calibration. Survivor bias can affect predictions at later time points, as the biologically oldest individuals are no longer present. Stochastic age analysis was carried out as previously described by Meyer and Schumacher^16^. A maximum stochastic age was set at 16.

### *C. elegans* microscopy and image analysis

*C. elegans* worms were anesthetized by 50 mM sodium azide or 1 mM levamisole. Anesthetized worms were placed on glass slides with an agar pad. *C. elegans* muscle images were taken using Spinning Disc Confocal Andor Dragonfly at 20x or 40x magnification. A z-stack was taken for the thickness of the worm at 1 or 2 um increments. A maximum intensity projection was created using ImageJ. Muscle filaments and mitochondria were identified by Labkit segmentation thresholding. Individual muscle cells were outlined using the polygon selection tool, and analyzed by MitochondriaAnalyzer ImageJ plugin. HIL-1::GFP punctae were quantified by analyzing z-stack images with IMARIS spot analysis.

### RNAi in *C. elegans*

HT115 bacteria carrying RNAi plasmids were used from Ahringer and Vidal libraries for feeding-based RNAi. HT115 bacteria were cultured overnight in LB media containing 100 μg/mL ampicillin at 37°C with gentle shaking. For the RNAi screen, the overnight culture was used directly on RNAi plates. All other RNAi experiments were carried out by using the overnight culture to inoculate a secondary culture until OD 0.6-1.0, and then concentrated 5-fold before applying and spreading on RNAi plates. The sequence of the RNAi plasmid was checked by Sanger sequencing to confirm that the insert targets the intended target gene for positive hits in the RNAi screen. HT115 bacteria containing an RNAi plasmid targeted towards *luciferase* was used as a negative control (*luci* ctrl.).

### Motility measurements in *C. elegans*

To determine motility, individual worms were transferred to M9 buffer, and the number of body bends in either direction were counted during 20 s. Worms that did not show any thrashing behavior were excluded from the analysis. Motility measurements were done in the *ergo-1(gg98)* mutant background to enhance the RNAi effect in neuromuscular tissues. *ergo-1(gg98)* mutant worms were healthier at the 5 w time point compared to WT worms, so a later time point (6 w) was used for the long-term ARD recovery experiments.

### CUT&RUN chromatin profiling

CUT&RUN was carried out and analyzed based on a protocol by Emerson and Lee^43^. In brief, N2 and *hil-1*::*gfp*::*flag* worms were induced into ARD as described above. After 1 week of ARD, worms were washed off ARD plates, combining 2-3 ARD plates (3000-4000 worms) for one sample. The washing steps of the original protocol were omitted as ARD worms are clear of bacteria. Worms were dissociated by incubation in SDS- and DTT-containing Cuticle Disruption Buffer followed by douncing in Wash Buffer, using a tissue grinder. The resulting mixture of worm chunks and cells was bound to activated Concanavalin A-coated beads. The respective primary antibodies (anti-FLAG (cat. no. 1804, Sigma Aldrich) and anti-H3 (cat. no. 1791, Abcam)) were added at a final concentration of 1:200 in Antibody Buffer, and samples were incubated rotating overnight at 4°C. On the following day, samples were washed and then incubated with Protein-A-Protein-G-MNase (pAG-MNase) which binds to the target antibody. MNase-digestion of the DNA bound by the target antibody was initiated through addition of calcium at 0°C. The reaction was stopped and the cleaved DNA fragments were released through sample incubation at 37°C. The DNA fragments were isolated, and library preparation for high-throughput sequencing was performed using the NEBNext Ultra II DNA Library Prep Kit for Illumina (cat. no. E7645S, New England Biolabs). Half of DNA obtained through CUT&RUN was used for library preparation. Size selection and purification were performed using Ampure XP beads, as stated. Evaluation of size distribution of the libraries was performed using a TapeStation. Samples were pooled at equal molarity (20 nM) and libraries were sequenced on Illumina NovaSeq (SP) by the Cologne Center for Genomics, with 20 million paired end reads (2x50 bp) per sample. Reads were aligned using STAR (2.7.10b) aligner with --outSAMtype BAM SortedByCoordinate --alignIntronMax 1 options to the WBcel235/ce11 genome to output a bam file. Bam index files were created using samtools view command. Indexed bam files were combined and used for broad peak calling with MACS3^66^ using --f BAM --g ce -- call summits - B --q 0.01 --keep-dup. Four biological replicate samples of *hil-1*::*gfp*::*flag* were compared against antibody background (FLAG antibody in WT background). Duplicate reads were kept as recommended for CUT&RUN by Emerson and Lee^43^. Peak profiles relative to transcription start/end sites, TEs, and intron-exon boundaries were produced by first creating a matrix file by computeMatrix, then plotting differential enrichment using plotHeatmap on the Galaxy server.

### Cell culture and treatments

All cell lines were grown at 37°C and 5% CO_2_. Human female embryonic kidney HEK293FT cells (cat. no. R70007, Invitrogen; Research Resource Identifier (RRID): CVCL_6911) were cultured in high-glucose Dulbecco’s Modified Eagle Medium (DMEM) (cat no. 41965039,

Gibco) supplemented with 10% fetal bovine serum (FBS) (cat. no. F7524, Sigma Aldrich, or cat. no. S1810, Biowest). Human male diploid lung WI-26 SV40 fibroblasts (WI-26 cells; cat. no. CCL-95.1, ATCC; RRID: CVCL_2758) were cultured in DMEM/F12 GlutaMAX medium (cat. no. 31331093, Thermo Fisher Scientific) containing 10% FBS. Both media were supplemented with 1% Penicillin–Streptomycin (cat. no. 15140122, Gibco). Both cell lines were confirmed to be *Mycoplasma*-free by regular testing for *Mycoplasma* contamination, using a PCR-based approach.

Amino acid (AA) starvation experiments were performed as previously described^67,68^. For WI-26 cells, DMEM/F12 GlutaMAX free of AAs (cat. no. D9811-01, US Biological) was used while starvation medium for HEK293FT cells was formulated according to the Gibco recipe for high glucose DMEM, omitting all AAs. The AA-free high glucose DMEM was filtered through a 0.22 μm filter device and tested for proper pH and osmolarity before use. For the respective AA-replete treatment media, high-glucose DMEM or DMEM/F12 GlutaMAX was used. Both starvation (-AA) and basal (+AA) treatment media were supplemented with 10% dialyzed FBS (dFBS) and 1% Penicillin–Streptomycin. For this purpose, FBS was dialysed against 1× PBS through 3500 MWCO dialysis tubing. For basal and starvation conditions, the culture medium was replaced with +AA and -AA treatment medium, respectively, 1 d before lysis. Addback was performed by replacing -AA medium with +AA medium after the indicated times. For inhibition of mTOR, the ATP-competitive inhibitor Torin1 (cat. no. 4247, Tocris) was dissolved in DMSO and added to freshly prepared full media to attain a final concentration of 250 nM. Treatment with the histone deacetylase inhibitor Vorinostat (cat. no. S1047, Selleckchem) was performed similarly to attain a final concentration of 10 µM. For experiments involving AA addback, siRNA-mediated gene silencing and for transcriptomics experiments, cells were seeded onto plates coated with Poly-L-Lysine (PLL) solution (cat no. sc-286689, Santa Cruz Biotechnology). PLL-coated coverslips were used for immunofluorescence experiments.

### Antibodies for cell culture experiments

Antibodies against phospho-p70 S6K (Thr389) (cat. no. 97596), total p70 S6K (cat. no. 34475), H3 (cat. no. 9715), Acetyl-K (cat. no. 9814), FOXO1 (cat. no. 2880) and FOXO3a (cat. no. 12829) were purchased from Cell Signaling Technology. A monoclonal antibody against H1.0 was purchased from Sigma Aldrich (05-629-I). An antibody against H1.2 was purchased from Covalab (pab0653).

### Generation of CRISPR knockout cell line

The HEK293FT *H1-0* knockout (KO) cell line was generated by gene editing using the pX459-based CRISPR/Cas9 system, as described elsewhere ^69^. For generating the single guide RNA (sgRNA) expression vectors, the DNA oligonucleotides were cloned in the BpiI restriction sites of the pX459 vector (cat. no. 62988, Addgene). The oligo sequences used for the sgRNA-expressing plasmids are provided in Extended Data Table 37. In brief, transfected cells were selected with 3 µg/mL puromycin (cat. no. A11138-03, Thermo Fisher Scientific) 31 h post transfection. Single clones were obtained by single cell sorting. KO clones were then first validated by immunoblotting and the mutation was subsequently confirmed by TOPO-TA cloning and DNA-sequencing. Negative control cell lines were created using an empty pX459 vector and the same procedure as described above for the *H1-0* KO cell line.

### Live-cell imaging

For live-cell imaging experiments, cells were seeded onto PLL-coated plates. On the subsequent day, culture media was replaced with +AA/-AA media and plates were immediately placed into the Incucyte to monitor cell viability and proliferation. For long-term starvation treatments, cells were deprived of AAs for 10 d. For AA addback experiments, cells were deprived of AAs for 1 d or 5 d, followed by addback of all AAs. Cell death was measured using the Incucyte Cytotox Dye (cat. no. 4632, Sartorius) according to the manufacturer’s instructions.

### Gene silencing experiments in HEK293FT cells

For transient knockdowns, cells were transfected with a pool of four siGENOME small interfering RNAs (siRNAs; Horizon Discovery). An siRNA duplex targeting *Renilla reniformis luciferase* (cat. no. P-002070-01-50, Horizon Discoveries) was used as a control. The oligonucleotide sequences are provided in Extended Data Table 37. Transfections were performed with 20 nM siRNA and Lipofectamine RNAiMax transfection reagent (cat. no. 13778075, Thermo Fisher Scientific), according to the manufacturer’s instructions. Treatments for starvation experiments were initiated 2 d after transfection by replacing media with +AA/– AA treatment media (free of siRNA). Cells were collected 3 d post-transfection and knockdown efficiency was verified by immunoblotting.

### mRNA isolation from HEK293FT cells and cDNA synthesis

Total RNA was isolated using the RNeasy Mini Kit (cat. no. 74106, Quiagen) according to the manufacturer’s instructions. For cDNA synthesis, 1 µg mRNA was reverse transcribed using the RevertAid H minus first strand cDNA synthesis kit (cat. no. K1631, Thermo Fisher Scientific) and an oligodT primer according to the manufacturer’s instructions.

### Real-time quantitative PCR

For real-time quantitative PCR (RT-qPCR) experiments, cDNA was diluted 1:50 in nuclease-free water. 4 µL diluted cDNA were mixed with 5 µL 2× qPCR SYBRGreen master mix with ROX (cat. no. K0223, Thermo Fisher Scientific) and 1 µL primer mix (2.5 µM of each primer). Technical triplicates were run and relative gene expression was calculated using the delta-delta-Ct method. *CNOT4* was used as a housekeeping control. The primer sequences are provided in Extended Data Table 37.

### Immunoblotting

For direct lysis in Laemmli buffer, cells were collected in serum-free DMEM to remove FBS and lysed by addition of Laemmli lysis buffer (100 mM NaF, 2 mM NaV, 0.011 g ml^-1^ β-glycerophosphate, 1× PhosSTOP phosphatase inhibitors (cat. no. 4906837001, Roche), 1× cOmplete protease inhibitors (cat. no. 11836153001, Roche), 1x Laemmli (6x Laemmli buffer: 350 mM Tris-HCl pH 6.8, 30% glycerol, 600 mM dithiothreitol, 12.8% sodium dodecyl sulfate (SDS), 0.12% bromphenolblue)) to the cell pellet. After resuspension in lysis buffer, samples were immediately moved to 95°C for 5 min. Once cooled down, each of the denatured samples was treated 10 µL Benzonase mix (1.95 µL H_2_O, 5.32 µL 1 M Tris-HCl pH 7.5, 1.4 µL 1M MgCl_2_ and 1.33 µL Benzonase (10.000 units/mL; cat. no. E1014-5KU, Sigma Aldrich)). Samples were incubated for 15 min at 37°C, 1500 rpm. Total proteins extracted by direct lysis were subjected to electrophoretic separation by SDS-polyacrylamide gel electrophoresis (SDS-PAGE) and analyzed by standard immunoblotting techniques. In brief, samples were transferred to 0.2 µm nitrocellulose membranes (cat. no. 10600002, Amersham) and stained with 0.2% Ponceau S solution (cat. no. 33427.01, Serva) to confirm equal loading. Normalization was performed by Ponceau S staining, normalizing to total protein levels by quantification with the GelAnalyzer software (v19.1; www.gelanalyzer.com). Membranes were blocked in 5% powdered milk (cat. no. 42590, Serva) in PBS-T (1× PBS and 0.1% Tween-20 (cat. no. A1389, AppliChem)) for 1 h at room temperature, washed 3 times for 5 min with PBS-T and incubated with primary antibodies in 5% BSA (cat. no. 10735086001, Roche), rotating overnight at 4 °C. All antibodies were used 1:1,000, apart from H3 (1:10,000). For H3, half the amount of protein was used for immunoblotting and primary antibody incubation was performed for 1 h at room temperature. After primary antibody incubation, membranes were washed 3 times for 5 min in PBS-T and incubated with the respective HRP-conjugated secondary antibodies (1:10,000 in PBS-T, 5% milk) for 1.5 h at room temperature. Signals were detected by enhanced chemiluminescence, using the ECL Western Blotting Substrate (cat. no. W1015, Promega). Immunoblot images were captured on film (cat. no. 47410-19284, Fuji) and quantified using the GelAnalyzer software.

### Immunofluorescence

For immunofluorescence experiments, HEK293FT cells were seeded onto PLL-coated glass coverslips and treated on the subsequent day as indicated above. After treatment, cells were fixed in the well with 4% Methanol-free Formaldehyde (cat. no. 28908, Thermo Fisher Scientific) in PBS for 10 min at room temperature. Samples were permeabilized with PBS-T solution (1x PBS and 0.1% Tween-20) in subsequent washing steps performed on 50 µL drops inside a humified chamber. After 2 PBS-T washes for 10 min at room temperature, samples were blocked with BBT solution (1x PBS, 0.1% Tween-20, 0.1% BSA) for 45 min at room temperature. Stainings were performed with H1.0 antibody, diluted 1:300 in BBT solution, overnight at 4°C. After 4 washes with BBT solution for 10 min each, samples were incubated with highly cross-absorbed secondary fluorescent anti-mouse Alexa Fluor-conjugated antibody (cat. no. 115-545-003, Jackson ImmunoResearch), diluted 1:500, for 1 h at room temperature. Samples were washed 2 times with PBS-T for 15 min each before staining with 4,6-diamidino-2-phenylindole (DAPI; cat. no. A1001, VWR), diluted 1:1500 in PBS-T, for 10 min at room temperature. After a last 15 min PBS-T wash at room temperature samples were mounted on slides using Fluoromount-G medium (cat. no. 00-4958-02, Thermo Fisher Scientific). Slides were dried protected from light at least overnight before imaging. Images were captured using a Spinning Disc Confocal Andor Dragonfly at 40x magnification.

### HEK293FT cells RNA-sequencing and transcriptome analysis

Gene expression changes were analyzed in an RNA-seq experiment. To this end, total RNA was isolated from HEK293FT *H1-0* KO cells and WT cells after 1 d +AA/-AA treatment using the RNeasy Plus Mini Kit (cat. no. 74134, Qiagen) and QIAshredder columns (cat. no. 79654, Qiagen), according to the manufacturer’s instructions. All experiments were performed in quadruplicates. Library preparation was done according to the Illumina TruSeq ribo zero (gold) protocol, with addition of ERCC RNA spike-ins. Sequencing-by-synthesis was performed on Illumina HiSeq by the Cologne Center for Genomics, with 50 million reads per sample, using a paired end 2x100 bp protocol. RNA-seq reads were analyzed through a standardized pipeline established by the MPI-AGE bioinformatics facility. RNA-seq reads were aligned to the Ensembl Homo sapiens release 105/ hg38 reference genome using kallisto^62^, and pair-wise differential gene expression was calculated using DESeq2^47^.

### Transcriptome interaction analysis (HEK293FT and *C. elegans*)

To determine which transcriptional changes occur linker histone-dependently during AA starvation and ARD/ARD recovery, we performed genotype-diet-interaction analyses. DESeq2 was used to model the effects of genotype and diet, as well as their interaction. First, transcript-level counts were aggregated into gene-level counts using tximport. A generalized linear model was then fitted to the count data, using the formula

∼ genotype + diet + genotype:diet,

where genotype represents the genetic background (i.e. WT or mutant), diet refers to the nutritional state/treatment condition (i.e. fed/basal conditions or ARD/ARD recovery/AA starvation), and genotype:diet is the interaction term. To account for differences in sequencing depth and RNA composition across samples, counts were normalized through DESeq2’s size factor normalization. Genes with low expression (total counts < 10) across all samples were excluded from the analysis. To improve the resulting estimated fold-changes, log_2_ fold shrinkage was applied using the apeglm method^35^.

For downstream analyses, significant genes that emerged from the interaction analysis (Adj. *P* value <0.05) were filtered for diet-responsive genes, i.e. genes for which expression was significantly changed (Adj. *P* value <0.05) either positively or negatively in response to the diet in the genotype control (i.e. N2 worms or WT cells). Then, the genes were grouped into 4 categories based on the direction of their regulation in response to the diet (upregulation vs. downregulation) and the effect of the gene perturbation (inhibited response vs. amplified response).

### Killifish fasting-refeeding and RNA-sequencing

African turquoise killifish *Nothobranchius furzeri* GRZ-AD strain was used for experiments. All killifish experiments were approved by Landesamt für Natur, Umwelt und Verbraucherschutz Nordrhein Westfahlen, Permit # AZ 81-02.04.2019.A055. Killifish maintenance and adipose tissue extraction for RNA-seq was carried out as previously described ^18^. Briefly, adipose tissues were extracted and washed twice with 1x PBS, and incubated with 500 µL dissociation mix (0.25% trypsin-EDTA (cat. no. 25200-056, Gibco), collagenase (4 mg/mL; cat. no. C9891, Sigma Aldrich) and DNase (100 ng/mL; cat. no. DN25-10MG, Sigma Aldrich). Adipose samples were incubated at 30°C for 45 min and pipetted until homogenized, and 800 µL DMEM-10% FBS were added, and centrifuged for 5 min at 400 r.c.f. Pellets following centrifugation were washed twice with 800 µL DMEM-10% FBS and resuspended in 500 µL XF Base Medium Minimal DMEM (Aglient Technologies), then filtered through a 70 µm nylon mesh and used for RNA extraction with RNeasy Mini kit (QIAGEN).

### Patient population and extraction of human PBMCs

Human PBMCs were obtained from male patients performing a 3 d water fast who participated in the RESET-PKD trial, a pilot study examining the impact of fasting in autosomal dominant polycystic kidney disease (ADPKD). Patients with ADPKD aged 18-60 years, who were otherwise considered healthy, were included in this non-randomized, non-blinded, single-center study at the University Hospital Cologne, described in detail elsewhere^70^. Heparinized whole blood was obtained from study participants after obtaining written informed consent and all study procedures were approved by the ethics committee of the Medical Faculty at the University of Cologne, Cologne, Germany (20-1040). The study was conducted in accordance with the Declaration of Helsinki and the good clinical practice guidelines by the International Conference on Harmonization and is registered on www.clinicaltrials.gov (NCT04472624).

Briefly, patients underwent complete water fasting for 3 d or maintained their usual dietary habits. For each patient, blood was drawn twice prior to the intervention (i.e. on *ad libitum* diet for all patients) to maintain baseline levels and once at the very end of the intervention, i.e. either while still fasting or maintaining their usual dietary habits. The time span between the first 2 visits ranged between 13 and 28 days, while the intervention was always started 7 days after the second initial blood draw. All patients presented in this study adhered to their prescribed dietary regimen and were still in ketosis at the end of the intervention.

PBMCs were extracted using ficoll density gradient centrifugation as follows: roughly 8 ml of heparinized whole blood were diluted 1:1 with PBS in sterile conditions and transferred to a Leucosep tube (cat no. GREI163290_500, Greiner Bio-One). Following centrifugation, PBMCs became visible as a white ring in the plasma fraction of the sample and were collected by manual aspiration. PBMCs were then counted and assessed for viability using Trypan blue staining in a semi-automated fashion employing the Countess® II FL Automated Cell Counter (Thermo Fisher). Subsequently, PBMCs were aliquoted, so that 1 × 10^6^ viable PBMCs were resuspended in 1 ml FBS +10% DMSO and cryopreserved in liquid nitrogen.

### RNA extraction from human PBMCs and RT-qPCR

Aliquots of 1 × 10^6^ viable PBMCs were thawed on ice. RNA was extracted employing a column based approach, using the RNeasy Micro Kit from (cat. no. 74004, Qiagen) as per the manufacturer’s instructions. To ensure complete removal of any remaining genomic DNA, an additional treatment with DNAse was performed using the RapidOut DNA Removal Kit (cat. no. K2981, Thermo Fisher Scientific) according to the manufacturer’s instructions. The concentration and purity of the RNA were measured on a NanoPhotometer® (Implen, NP80). RT-qPCR was run in technical quadruplicates with *RPLP0* as a housekeeping control. Relative gene expression was calculated using the delta-delta-Ct method. The primer sequences are provided in Extended Data Table 37.

## Data availability

Raw sequencing data will be available upon publication on SRA or PRIDE repositories. Worm strains are available upon request. Processed data are available in the main text or supplementary materials.

## Acknowledgments

We thank Felicity Emerson for discussion and advice about CUT&RUN. We thank the MPI-AGE bioinformatics core facility, especially Ayesha Iqbal, Yun Wang, and Jorge Boucas. We thank the MPI-AGE FACS & Imaging core facility at MPI-AGE, especially Marcel Kirchner. We thank the Cologne Center for Genomics for RNA and DNA sample sequencing. We thank Antebi lab members for continued discussion and input on our study. We thank Sarah Kreuz, Wenming Huang, Tim Nonninger, Youngjun Park and Victoria Martinez-Miguel for constructive feedback on our manuscript. We thank Nadine Hochhard, Anna Löhrke, Luca Jeromin, Tim Droth and Rosa Wärner for technical assistance. We received worm strains from CGC and NBRP. Some figure panels and graphical abstract were created using BioRender.

## Funding

Max Planck Society (AA, CD)

Cologne Graduate School of Ageing (ARD, FA, DS, DHM)

Deutsche Forschungsgemeinschaft (German Research Foundation) under Germany’s Excellence Strategy, CECAD; EXC 2030-390661388 (AA, CD, R-UM)

Deutsche Forschungsgemeinschaft (German Research Foundation) Research Unit Grant FOR2722 DE3170/1-1; Project No. 384170921 (CD)

European Research Council under the European Union’s Horizon 2020 research and innovation programme; 757729 (CD)

Japan Society for Promotion of Science Overseas fellowship; 202260346 (KK)

## Author contributions

Conceptualization: KK and AA conceived of the study.

Investigation: KK, ARD, BG, CL, DS carried out worm experiments. RR carried out killifish experiments. ARD, SF, FA carried out cell culture experiments. JDS, SO, FG carried out human patient experiments. KK, ARD, DHM carried out bioinformatic analyses.

Funding Acquisition: AA (worm, killifish, cell culture, human patient samples), CD (cell culture), R-UM (human patient samples).

Supervision: AA (worm, killifish, cell culture, human patient samples), CD (cell culture), R-U M (human patient samples).

Writing: KK, ARD, AA wrote the manuscript with input from all authors.

## Extended data figures and tables

**Extended Data Fig. 1.**
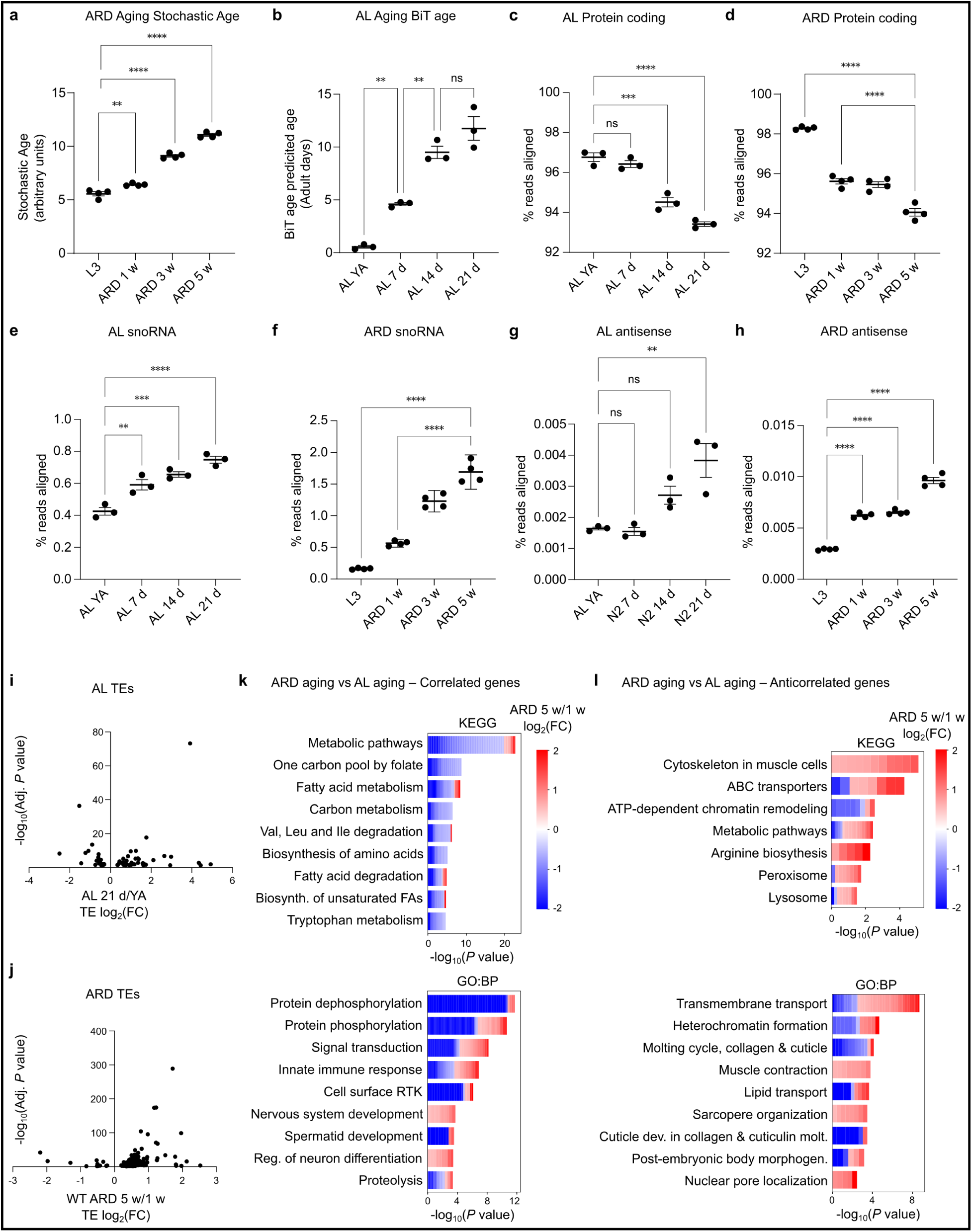
*Ad libitum* aging and adult reproductive diapause aging share similar transcriptomic changes. **a**, Stochastic age predictions based on transcriptomes from worms prior to ARD induction at mid-L3 stage, and ARD for 1, 3, and 5 weeks. **b**, BiT age quantifications of AL aging samples. Maximum biological age set at 15.5 days (see Methods-Age prediction analysis for details). **c**, Percentage of transcriptome reads aligned to protein coding regions from worms during AL aging (Young adult, 7 d, 14 d, 21 d), and **d**, during ARD. **e**, Percentage of transcriptome reads aligned to snoRNA from worms during AL aging, and **f**, during ARD. **g**, Percentage of transcriptome reads aligned to antisense RNA from worms during AL aging, and **h**, during ARD. **i**, Changes in TE expression during AL aging, and **j**, during ARD. **k**, Enriched KEGG and GO:BP gene ontology terms for differentially expressed genes (DEGs) with correlated, and **l**, anticorrelated gene expression during AL aging and ARD aging. One-way ANOVA with multiple comparisons \**P* < 0.05; \*\**P* < 0.01, *P*\*\*\* < 0.001. Error bars represent mean ± SEM.

**Extended Data Fig. 2.**
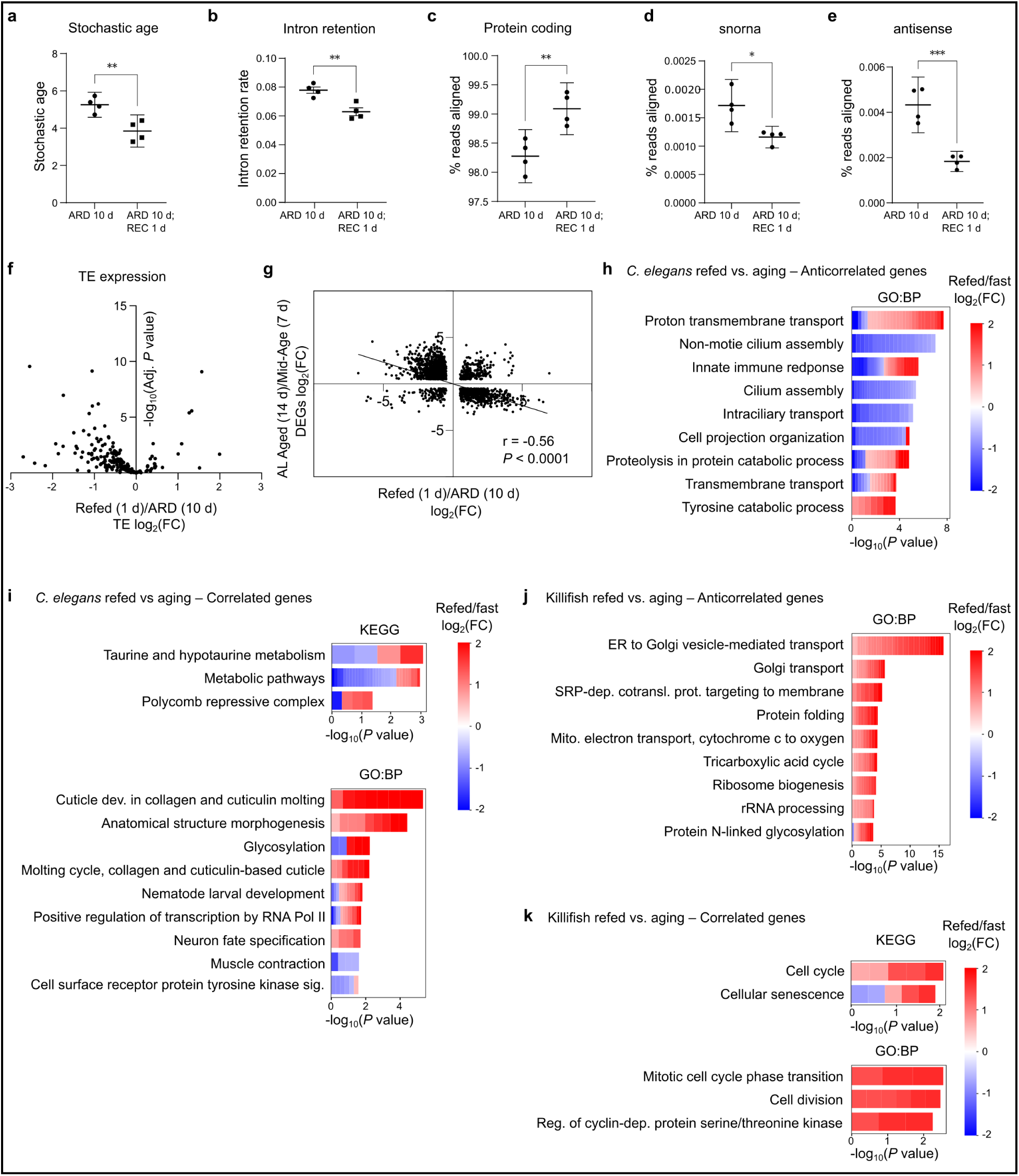
Refeeding reverses age-related transcriptome changes. **a**, Stochastic age predictions based on transcriptomes from worms in ARD 10 d and Refed 1d. **b**, Percentage of transcripts with intron retention in ARD and Refed samples. **c**, Percentage of transcriptome reads aligned to protein coding regions from ARD and Refed samples. **d**, Percentage of transcriptome reads aligned to snoRNA from ARD and Refed samples. **e**, Percentage of transcriptome reads aligned to antisense RNA from ARD and Refed samples. **f**, Volcano plot of TE expression from ARD and Refed samples. **g**, Correlation plot of AL aging (14 d vs 7 d) and Refed (1d) vs ARD (10 d); germline genes were removed to rule out that the inverse correlation occurs solely from re-expression of germline-related genes during refeeding from ARD. Different aging time points were used in this plot compared to Fig. 1E to assess a separate stage of aging. **h**, GO:BP terms for anticorrelated significant DEGs in AL aging (Day 14 vs YA) and Refeeding (Refed 1 d v ARD 10 d). **i**, KEGG and GO:BP terms for correlated DEGs in AL aging (Day 14 vs YA) and Refeeding (Refed 1 d vs ARD 10 d). **j**, KEGG and GO:BP terms for anticorrelated DEGs in killifish AL aging (18 w vs 7 w) and Refeeding (Refed 1 d vs Fast 3 d). **k**, KEGG and GO:BP terms for correlated DEGs in killifish AL aging (18 w vs 7 w) and Refeeding (Refed 1 d vs Fast 3 d). Unpaired t-test \**P* < 0.05; \*\**P* < 0.01, *P*\*\*\* < 0.001. Error bars represent mean ± SEM.

**Extended Data Fig. 3.**
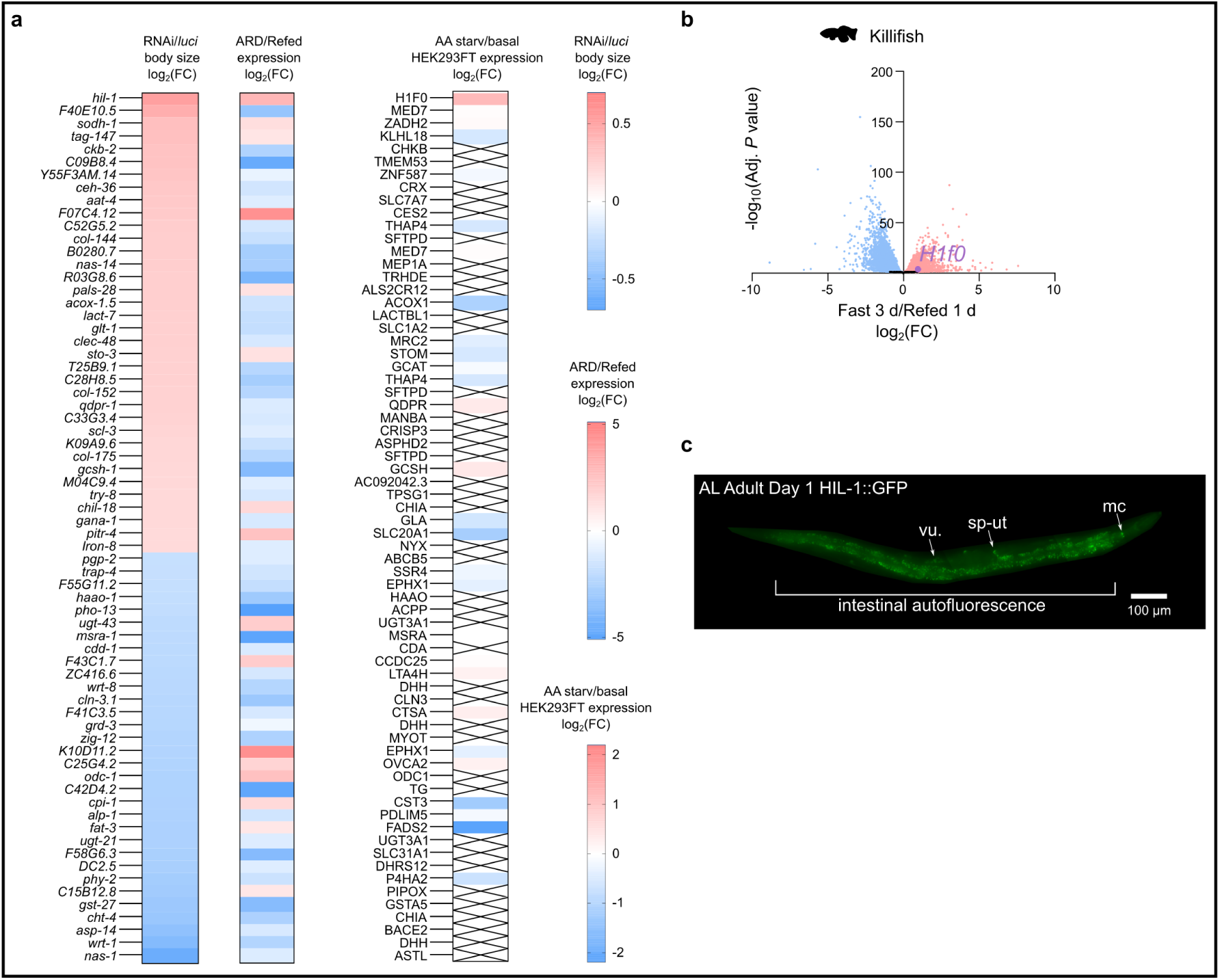
*hil-1/H1-0* is an evolutionarily conserved nutrient-regulated linker histone. **a**, Heat map of body size effects in RNAi screen (first column). Heat map of gene expression changes in ARD 10 d vs. Refed 1 d (second column). Heat map of protein expression changes in –AA 1 d Starvation vs. +AA (AA-replete) in HEK293FT cells (third column). **b**, Volcano plot of DEGs from fasting (3 d) and refeeding (1 d) in killifish visceral adipose tissue. **c**, Fluorescence microscopy of an AL adult Day 1 *C. elegans* worm expressing endogenously tagged HIL-1::GFP.

**Extended Data Fig. 4.**
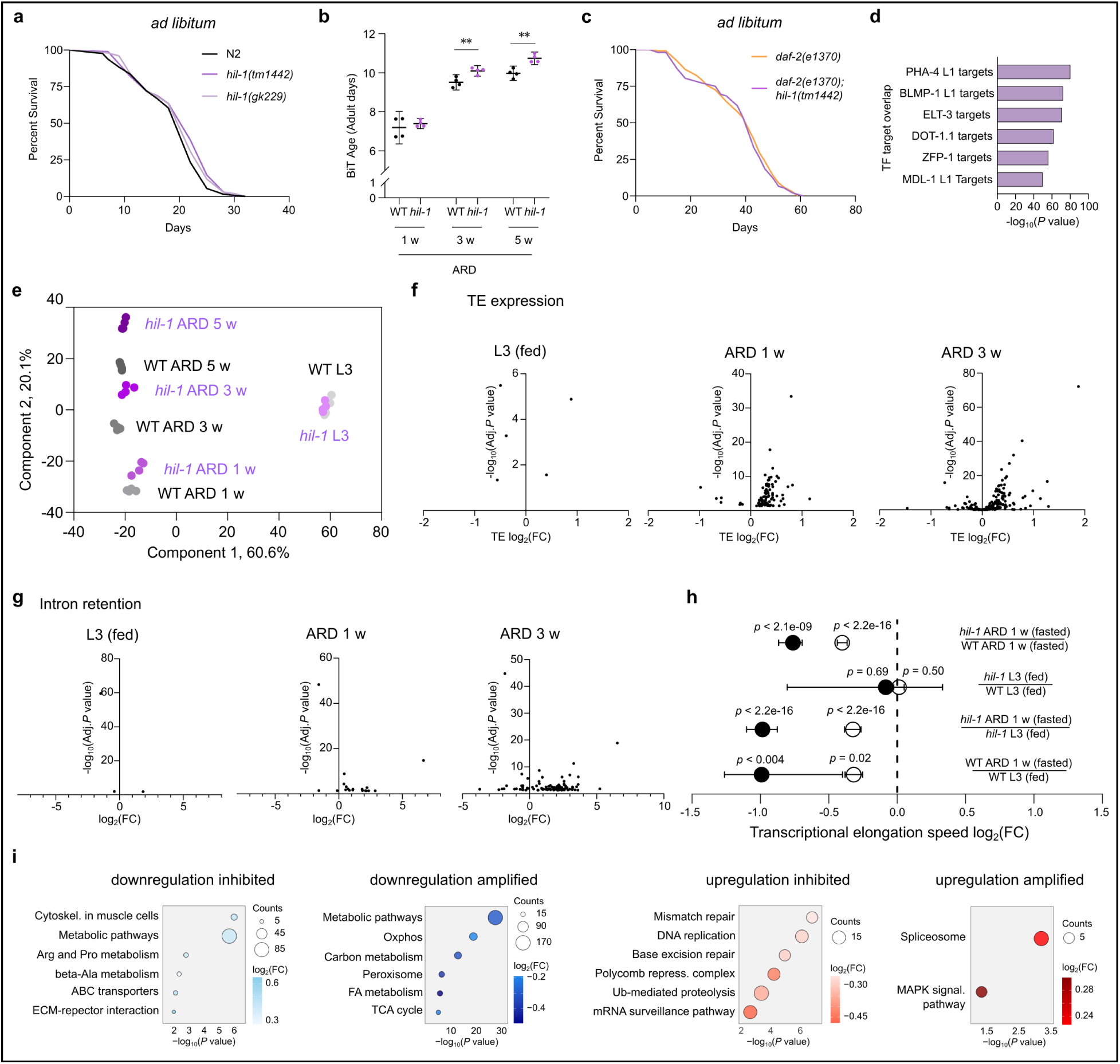
HIL-1 directly affects the transcriptome for fasting adaptation. **a**, AL lifespan of WT, *hil-1(tm1442)*, *hil-1(gk229)* worms. N = 3. **b**, BiT age biological age predictions based on transcriptomes from worms at ARD 1 w, ARD 3 w, and ARD 5 w in WT and *hil-1(tm1442)* mutant worms. **c**, AL lifespan of *daf-2(e1370)* and *daf-2(e1370); hil-1(tm1442)* worms. N = 3. Error bars represent mean ± SEM. **d**, Transcription factor overlap of HIL-1 direct binding targets using WormExp ‘TF Targets’. **e**, PCA plot of transcriptomes from worms at ARD 1 w, ARD 3 w, and ARD 5 w in WT and *hil-1(tm1442)* mutant worms. **f**, Volcano plots of differential expression of TE expression (*hil-1(tm1442)* vs. WT) in mid-L3, ARD 1 w, ARD 3 w worms. **g**, Volcano plots of differential intron retention of transcripts (*hil-1(tm1442)* vs. WT) in mid-L3, ARD 1 w, ARD 3 w worms. **h**, Transcriptional speed estimates of ARD 1 w *hil-1(tm1442)* vs. WT; mid-L3 *hil-1(tm1442)* vs. WT; ARD 1 w *hil-1(tm1442)* vs. mid-L3 *hil-1(tm1442)*; ARD 1 w WT vs. mid-L3 WT. Empty circles indicate results using all intronic reads. Solid circles indicate results using intronic reads with consistent effects across replicates. Error bars represent 95% confidence intervals. **i**, Top 6 enriched KEGG terms of significant genes from the genotype-ARD interaction analysis are displayed for each category.

**Extended Data Fig. 5.**
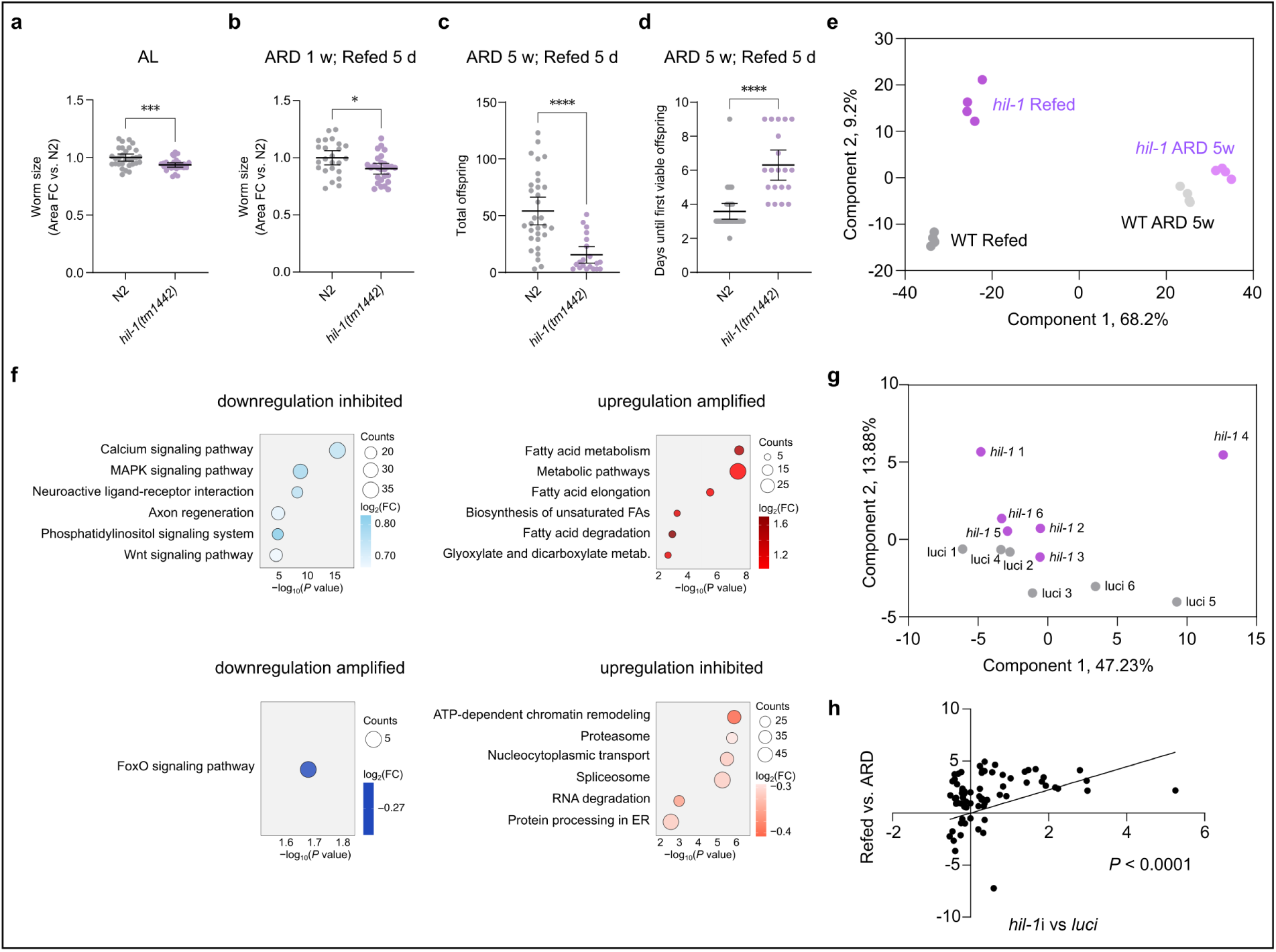
HIL-1 regulation is critical for refeeding-induced recovery. **a**, Body size of WT and *hil-1(tm1442)* in AL conditions measured 5 d after egg-lay, or **b**, Refed 5 d after ARD 1 w. **c**, Total offspring of WT and *hil-1(tm1442)* worms refed from ARD 5 w. **d**, Days until first viable offspring of WT and *hil-1(tm1442)* worms refed from ARD 5 w. **e**, PCA plot of transcriptomes of WT and *hil-1(tm1442)* worms from ARD 5 w and subsequently refed 2 d. **f**, Top 6 enriched KEGG terms of significant genes from the genotype-refeeding interaction analysis are displayed for each category. **g**, PCA plot of transcriptomes of WT worms 3 d after recovery from 5 w in ARD on *luci* ctrl. or *hil-1* RNAi. **h**, Correlation plot of significantly regulated genes in *hil-1* knockdown vs. *luci* ctrl. compared with Refed 1 d vs. ARD 10 d. Error bars represent mean ± SEM.

**Extended Data Fig. 6.**
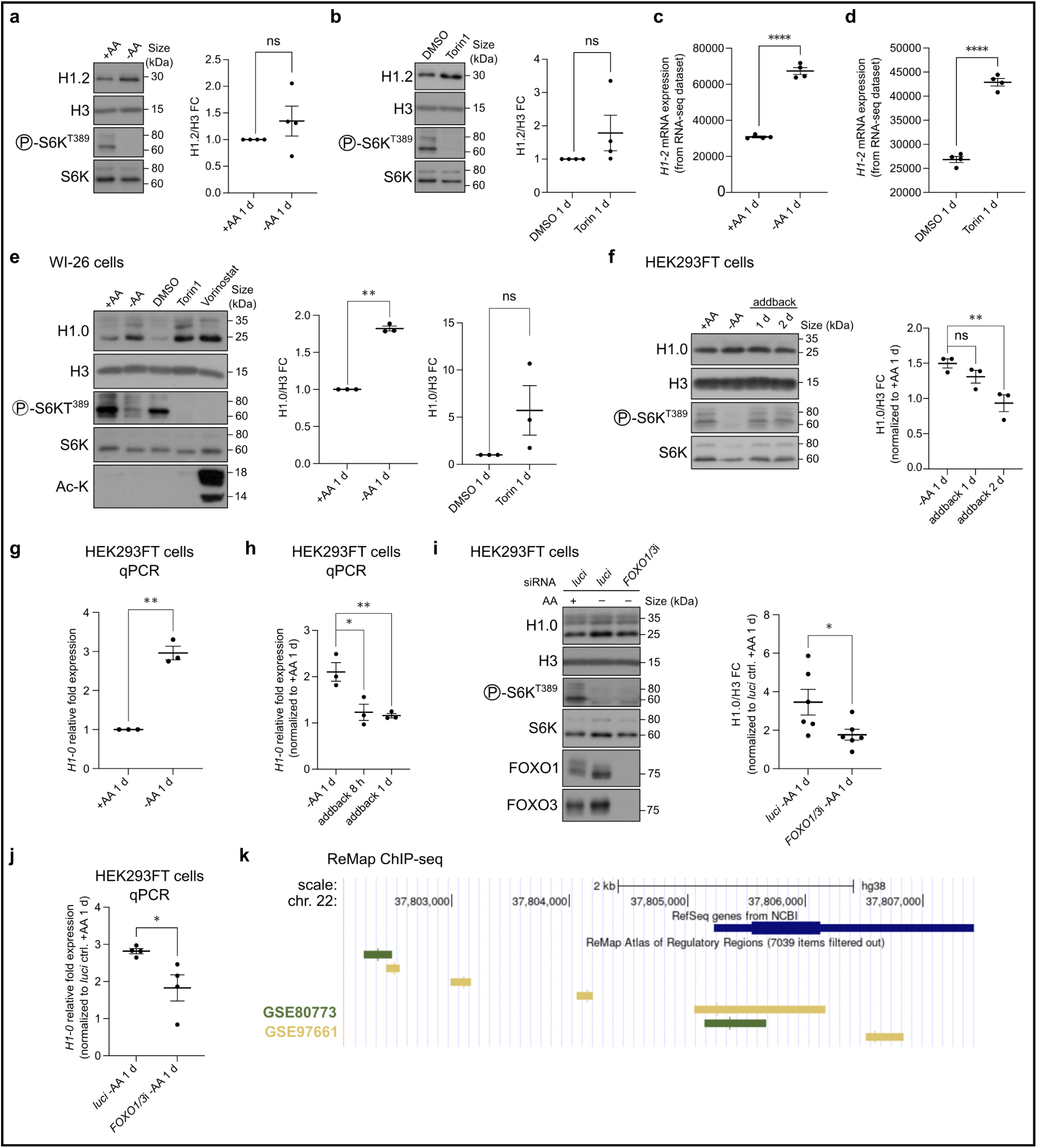
Evolutionarily conserved regulation of *hil-1/H1-0*. **a**, Immunoblots of lysates from HEK293FT cells under -AA 1 d Starvation vs. +AA 1 d (AA-replete) conditions, probed with the indicated antibodies. N = 4. **b**, Immunoblots of lysates from HEK293FT cells under Torin1 1 d (250 nM) vs. DMSO 1 d control conditions, probed with the indicated antibodies. N=4. **c**, RNA-seq expression of *H1-2* in HEK293FT cells under -AA 1 d Starvation vs. +AA 1 d. N = 4. **d**, RNA-seq expression of *H1-2* in HEK293FT cells under Torin1 1 d (250 nM) vs. DMSO 1 d control conditions. N = 4. **e**, Immunoblots of lysates from WI-26 cells under -AA 1 d Starvation vs. +AA 1 d, and Torin1 1 d (250 nM) vs. DMSO 1 d. Vorinostat (10 µM) was used as a control for H1.0 induction ^60^. N = 3. **f**, Immunoblots of lysates from HEK293FT cells under AA addback after -AA 1 d Starvation (addback for 1 d and 2 d) vs. -AA 1 d Starvation, probed with the indicated antibodies. Displayed data was normalized to AA-replete +AA 1 d condition. N = 3. **g**, *H1-0* mRNA levels were assessed by RT-qPCR in HEK293FT cells under -AA 1 d Starvation vs. +AA 1 d. Displayed data was normalized to +AA 1 d condition. CCR4-NOT transcription complex, subunit 4 (*CNOT4*) was used as a housekeeping gene. N = 3. **h**, *H1-0* mRNA levels were assessed by RT-qPCR in HEK293FT cells under AA addback after -AA 1 d Starvation (addback for 8 h and 1 d) vs. -AA 1 d Starvation. Displayed data was normalized to +AA 1 d condition. CCR4-NOT transcription complex, subunit 4 (*CNOT4*) was used as a housekeeping gene. **i**, Immunoblots of lysates from HEK293FT cells under the indicated siRNA-mediated knockdown conditions, probed with the indicated antibodies. AA Starvation 1 d was initiated 2 d after transfection. *CNOT4* was used as a housekeeping gene. Displayed data was normalized to *luci* ctrl. +AA 1 d condition. N = 3. **j**, *H1-0* mRNA levels were assessed by RT-qPCR in HEK293FT cells under the indicated knockdown conditions. AA Starvation 1 d was initiated 2 d after transfection. *CNOT4* was used as a housekeeping gene. Displayed data was normalized to *luci* ctrl. +AA 1 d condition. N = 3. **k**, ReMap ChIP-seq track in the UCSC Genome Browser, showing experimentally validated binding of FOXO1 (green peak, GSE80773, CD34 cells) and FOXO3 (yellow peak, GSE97661, Hep-G2 cells) in the *H1-0* promoter region. Error bars represent mean ± SEM. Statistical significance between two groups was tested by unpaired t-test or one-sample t-test (comparing against a hypothetical value of one) while the one-way ANOVA was used for multiple comparisons. *P*\* < 0.05; *P*\*\* < 0.01; *P\*\*\** < 0.001; *P\*\*\*\** < 0.0001. ns = not significant.

**Extended Data Fig. 7.**
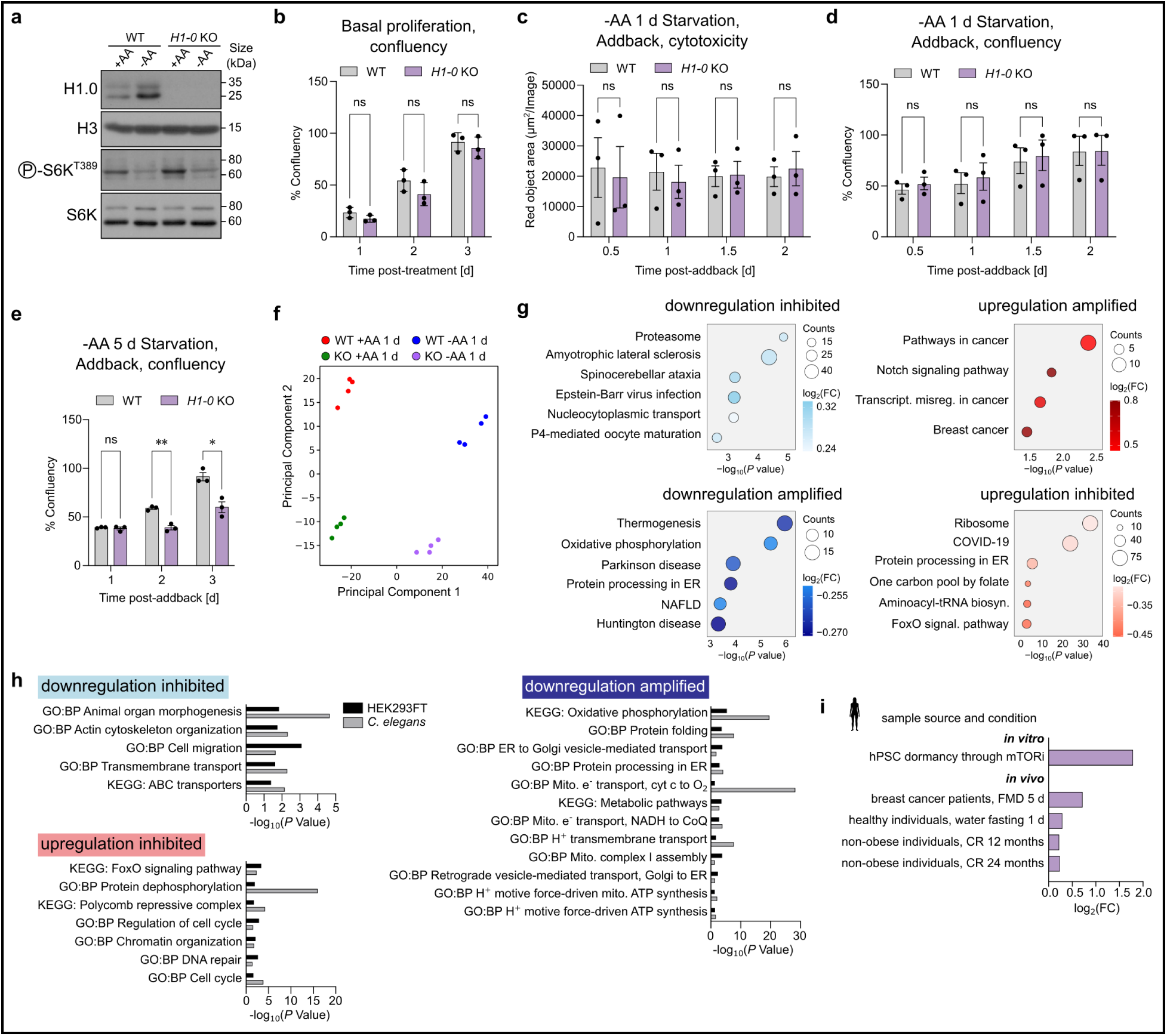
Evolutionarily conserved functions of *hil-1/H1-0*. **a,** Immunoblots of lysates from HEK293FT *H1-0* KO cells and WT cells under -AA 1 d starvation vs. +AA 1 d, probed with the indicated antibodies. **b**, Proliferation of *H1-0* KO cells under basal AA-replete conditions, as quantified by confluency over time. N = 3. **c**, Evaluation of cell death in HEK293FT *H1-0* KO cells upon AA addback after AA starvation 1 d, as measured by the Incucyte Cytotoxicity Assay. Cell death was quantified by Red object area. N = 3. **d**, Proliferation of *H1-0* KO cells upon AA addback after AA starvation 1 d, as quantified by confluency over time. N = 3. **e**, Proliferation of *H1-0* KO cells upon AA addback after AA starvation 5 d, as quantified by confluency over time. N = 3. **f**, PCA plot of transcriptomes of WT and *H1-0* KO cells under +AA 1 d condition and -AA 1 d starvation conditions. **g**, Top 6 enriched KEGG terms of significant genes from the genotype-starvation interaction analysis are displayed for each category. **h**, Overlapping KEGG and GO:BP terms from the interaction analyses of *hil-1* ARD 1 w worms and -AA 1 d Starved *H1-0* KO cells. Hypergeometric testing revealed significant enrichment of terms in the ‘up, inhibited’ category (*P* = 0.043) and in the ‘down, amplified’ category (*P* = 6.40^-10^). **i**, Summary of *H1-0* regulation from publicly available human *in vitro* and *in vivo* data across different low nutrient conditions. Error bars represent mean ± SEM. Statistical significance between genotypes in the Incucyte timecourse was assessed by multiple unpaired t tests with Holm-Šídák correction. *P*\* < 0.05; *P*\*\* < 0.01. ns = not significant.

**Extended Data Table 1.**

AL aging TE expression.

**Extended Data Table 2.**

ARD aging TE expression.

**Extended Data Table 3.**

DEGs during ARD aging and AL aging.

**Extended Data Table 4.**

GO:BP and KEGG term enrichments from DEGs in ARD and AL aging.

**Extended Data Table 5.**

Differentially expressed TEs during Refeeding and AL aging.

**Extended Data Table 6.**

DEGs during Refeeding and AL aging (14 d vs. 1 d).

**Extended Data Table 7.**

DEGs during Refeeding and AL aging (14 d vs. 1 d) with germline genes removed.

**Extended Data Table 8.**

DEGs during Refeeding and AL aging (14 d vs. 7 d) with germline genes removed.

**Extended Data Table 9.**

GO:BP and KEGG term enrichments from DEGs in Refeeding and AL aging.

**Extended Data Table 10.**

DEGs during Refeeding and AL aging in killifish.

**Extended Data Table 11.**

GO:BP and KEGG term enrichments from DEGs in Refeeding and AL aging in killifish.

**Extended Data Table 12.**

ARD REC REP transcriptomics normalized read counts.

**Extended Data Table 13.**

List of 380 genes for RNAi screen with body size effect.

**Extended Data Table 14.**

RNAi effect and expression changes of top candidates.

**Extended Data Table 15.**

DEGs from ARD (10 d)/Refed(1 d).

**Extended Data Table 16.**

DEGs from AA starvation (1 d)/basal in HEK293FT cells.

**Extended Data Table 17.**

DEGs from Torin1 (1 d)/DMSO in HEK293FT cells.

**Extended Data Table 18.**

DEGs from Fasted (3 d)/Refed (1 d) from killifish visceral adipose.

**Extended Data Table 19.**

Identified HIL-1 peaks.

**Extended Data Table 20.**

HIL-1 peaks overlapping with genes (including promoter and 3’UTR).

**Extended Data Table 21.**

HIL-1 target gene overlap with TF targets.

**Extended Data Table 22.**

Differential TE expression *hil-1(tm1442)*/WT.

**Extended Data Table 23.**

Differential intron retention in *hil-1(tm1442)*/WT.

**Extended Data Table 24.**

Genotype-ARD interaction analysis for *hil-1(tm1442)* mutant during ARD induction.

**Extended Data Table 25.**

GO:BP and KEGG enrichments of significant genes from genotype-ARD interaction analysis for *hil-1* mutant during ARD.

**Extended Data Table 26.**

Genotype-refeeding interaction analysis for *hil-1(tm1442)* mutant during refeeding (2 d) following ARD (5 w).

**Extended Data Table 27.**

GO:BP and KEGG enrichments of significant genes from genotype-refeeding interaction analysis for *hil-1* mutant during refeeding.

**Extended Data Table 28.**

DEGs from *hil-1* knockdown at 3 d refeeding following 5 w in ARD.

**Extended Data Table 29.**

GO:BP and KEGG enrichments of significant genes from *hil-1* knockdown at 3 d refeeding following 5 w in ARD.

**Extended Data Table 30.**

Genotype-starvation interaction analysis for *H1-0* KO cells during 1 d AA starvation.

**Extended Data Table 31.**

Differential TE expression -AA starvation 1 d *H1-0* KO/WT.

**Extended Data Table 32.**

Differential intron retention in -AA starvation 1 d *H1-0* KO/WT.

**Extended Data Table 33.**

GO:BP and KEGG enrichments of significant genes from genotype-starvation interaction analysis for *H1-0* KO during AA starvation.

**Extended Data Table 34.**

Overlap of GO:BP and KEGG enrichments from genotype-diet interaction analysis for *hil-1* ARD and *H1-0* KO AA starvation.

**Extended Data Table 35.**

*C. elegans* strain list.

**Extended Data Table 36.**

*C. elegans* lifespan experiment summary table.

**Extended Data Table 37.**

Oligonucleotide sequence list.

